# Coral reef restoration efforts in Latin American countries and territories

**DOI:** 10.1101/2020.02.16.950998

**Authors:** Elisa Bayraktarov, Anastazia T. Banaszak, Phanor Montoya Maya, Joanie Kleypas, Jesús E. Arias-González, Macarena Blanco, Johanna Calle Triviño, Nufar Charuvi, Camilo Cortés Useche, Victor Galván, Miguel A. García Salgado, Mariana Gnecco, Sergio D. Guendulain García, Edwin A. Hernández Delgado, José A. Marín Moraga, María Fernanda Maya, Sandra Mendoza Quiroz, Samantha Mercado Cervantes, Megan Morikawa, Gabriela Nava, Valeria Pizarro, Rita I. Sellares-Blasco, Samuel E. Suleimán Ramos, Tatiana Villalobos Cubero, María Villalpando, Sarah Frías-Torres

## Abstract

Coral reefs worldwide are degrading due to climate change, overfishing, pollution, coastal development, bleaching and diseases. In areas where natural recovery is negligible or protection through management interventions insufficient, active restoration becomes critical. The Reef Futures symposium in 2018 brought together over 400 reef restoration experts, businesses, and civil organizations, and galvanized them to save coral reefs through restoration or identify alternative solutions. The symposium highlighted that solutions and discoveries from long-term and ongoing coral reef restoration projects in Spanish-speaking countries in the Caribbean and Eastern Tropical Pacific were not well known internationally. Therefore, a meeting of scientists and practitioners working in these locations was held to compile the data on the extent of coral reef restoration efforts, advances and challenges. Here, we present unpublished data from 12 coral reef restoration case studies from five Latin American countries, describe their motivations and techniques used, and provide estimates on total annual project cost per unit area of reef intervened, spatial extent as well as project duration. We found that most projects used direct transplantation, the coral gardening method, micro-fragmentation or larval propagation, and aimed to optimize or scale-up restoration approaches (51%) or provide alternative, sustainable livelihood opportunities (15%) followed by promoting coral reef conservation stewardship and re-establishing a self-sustaining, functioning reef ecosystem (both 13%). Reasons for restoring coral reefs were mainly biotic and experimental (both 42%), followed by idealistic and pragmatic motivations (both 8%). The median annual total cost from all projects was $93,000 USD (range: $10,000 USD - $331,802 USD) (2018 dollars) and intervened a median spatial area of 1 ha (range: 0.06 ha - 8.39 ha). The median project duration was 3 years; however, projects have lasted up to 17 years. Project feasibility was high with a median of 0.7 (range: 0.5 - 0.8). This study closes the knowledge gap between academia and practitioners and overcomes the language barrier by providing the first comprehensive compilation of data from ongoing coral reef restoration efforts in Latin America.

## Introduction

Active restoration, the process of assisting the recovery of an ecosystem that has been degraded, damaged, or destroyed [1], may be increasingly necessary on coral reefs, once it has been determined that the natural recovery of corals is hindered [2]. The goal of any restoration action is to eventually establish self-sustaining, sexually reproducing populations with enough genetic variation enabling them to adapt to a changing environment [3–5].

Coral reef restoration may play a particularly important role where coral species are threatened with extinction. The Caribbean Elkhorn coral, *Acropora palmata*, and Staghorn coral, *A. cervicornis*, were once widely distributed and among the major reef-building species in the region [6]. Both species are now listed as Critically Endangered on the International Union for Conservation of Nature (IUCN) Red List [7] as a result of major losses in cover of both species throughout the Caribbean since the 1970s [8]. Management programmes have not aided in the recovery of *A. palmata* [9]. In this context, active restoration of these species is essential to recover their ecosystem functions in the Caribbean region.

Several techniques are used for the restoration of coral reefs. The most common techniques are based on asexual methods such as direct transplantation, coral gardening, and micro-fragmentation [10]. An alternative technique, larval propagation, is based on the collection of gametes and the consequent culturing of embryos and larvae, after which the coral spat are either grown in *ex situ* aquaria to largersized colonies or are outplanted onto degraded reefs at approximately one month old [11]. While the techniques used to restore coral reefs are reviewed elsewhere (e.g. [10, 12–14]), here we focus on direct transplantation, coral gardening, micro-fragmentation, and larval propagation as the techniques most-commonly employed by the case studies in the study area. One of the oldest techniques used in coral reef restoration is direct transplantation of corals [15], which involves the harvesting of coral colonies from a donor site and their immediate transplantation to a restoration site or re-attaching colonies that have been dislodged by a ship grounding, storm or hurricane [16]. The coral gardening approach was developed to scale-up restoration while reducing the stress on donor colonies. Fragments of corals are harvested from donor colonies, grown in nurseries to a threshold size [17] before being transplanted onto a degraded reef [18, 19]. Nurseries can be ocean-based (*in situ*) or land-based (*ex situ*). *In situ* nurseries are typically located at well-lit sites safe from predation, storm surges, and wave energy, and are regularly maintained and cleaned by physical removal of algal growth [20]. However, strategic siting of ocean nurseries can promote the recruitment of fish assemblages that eat biofouling, thus may significantly reduce person-hours spent in nursery cleaning [21]. *In situ* nurseries can have many shapes and sizes. For example, they can consist of floating mid-water structures built using ropes, mesh or cages [21–24], structures placed on concrete, tables or frames [25], PVC ‘trees’ [26], PVC grids or dead coral bommies [27]. *Ex situ* nurseries typically use flow-through large aquaria or raceways, and require continuous access to electricity, water quality monitoring, and control of temperature and light availability [28]. Micro-fragmentation is an approach especially useful for slow-growing massive corals. This technique involves the fragmentation of parts of a massive coral donor to yield multiple ~1 cm^2^ fragments. The fragments are placed close to each other on either artificial substrates or on the surface of dead coral colonies. The micro-fragments, as they recognize neighbouring fragments as kin, grow towards each other and fuse [29]. Ideally, they are outplanted to the degraded reef at a size of ~6 cm^2^ [29, 30]. Larval propagation involves the breeding of corals from eggs and sperm. Studies describing this technique typically report the use of raceways with seawater flow-through systems where coral spawn is collected from the wild, fertilization is assisted, embryos are cultured to larvae, which are settled onto substrates and then transported and seeded onto a degraded coral reef [31–33]. This process has also been referred to as larval enhancement, sexual propagation, sexual coral cultivation or larval reseeding [12]. As an emerging larval propagation technique, larval restoration concentrates coral larvae over enhancement plots on the degraded reef to facilitate coral larvae settlement directly to the substrate, without the need of laboratory facilities [34]. The main advantages of the larval propagation techniques are that they increase the genetic diversity among restored coral populations thus enabling increased rates of adaptation and improved resilience in the context of climate change [35], and they have the potential to be used over large scales while reducing the cost [31]. Also, they do not cause damage to the parent colonies.

While efforts in the USA, Australia or places where European scientists conduct their research are well described in the published literature and disseminated at conferences, there is a paucity of documentation on coral reef restoration projects carried out by practitioners in the Caribbean and Eastern Tropical Pacific. Reasons for this lack of exchange may be the language barrier, lack of interest in knowledge transfer between higher and lower income countries or cultural differences as well as lack of funding. In 2018, the Reef Futures symposium was held in the Florida Keys, USA and attended by over 400 delegates. The aim of this international meeting was to ‘bring together experts from around the world to share the latest science and techniques for coral reef restoration while kicking off a global effort to dramatically scale-up the impact and reach of restoration as a major tool for coral reef conservation and management’. The conference was organized by the Coral Restoration Consortium, which is ‘a community of practice comprised of scientists, managers, coral restoration practitioners, and educators dedicated to enabling coral reef ecosystems to survive the 21^st^ century and beyond’ [36]. Within the Reef Futures conference, we convened a meeting of scientists and practitioners involved in active coral reef restoration in the Latin- and Centro-American Caribbean as well as the Eastern Tropical Pacific to fill the knowledge gap between academia and practitioners in the region and overcome the language barriers in coral reef restoration. Here, we showcase the advances and share the lessons learned from 12 restoration case studies from the Caribbean and Eastern Tropical Pacific. We provide a comprehensive compilation of unpublished data from coral reef restoration efforts where we outline the techniques that were employed, the motivations and objectives of each project, total project cost per unit area per year, spatial extent of intervention, and project duration. This work provides the most complete data set on total project cost and feasibility of coral reef restoration from practical cases that may guide decisions required to establish new restoration projects in the future.

## Approach

### Data collection

The co-authors of this work contributed data and descriptions of their restoration projects which constitute the case studies used here. The coral reef restoration projects were carried out in Latin American countries and territories in the Caribbean and Eastern Tropical Pacific (**Fig. 1**). The data obtained included estimates on total annual project cost, spatial extent of area intervened, project duration, and an estimate on the project reaching specific objectives within a fixed period of time. The motivations for each restoration project were adopted from [10, 37, 38] and classified as biotic, experimental, idealistic, legislative, and pragmatic (**Table 1**).

**Figure 1:**
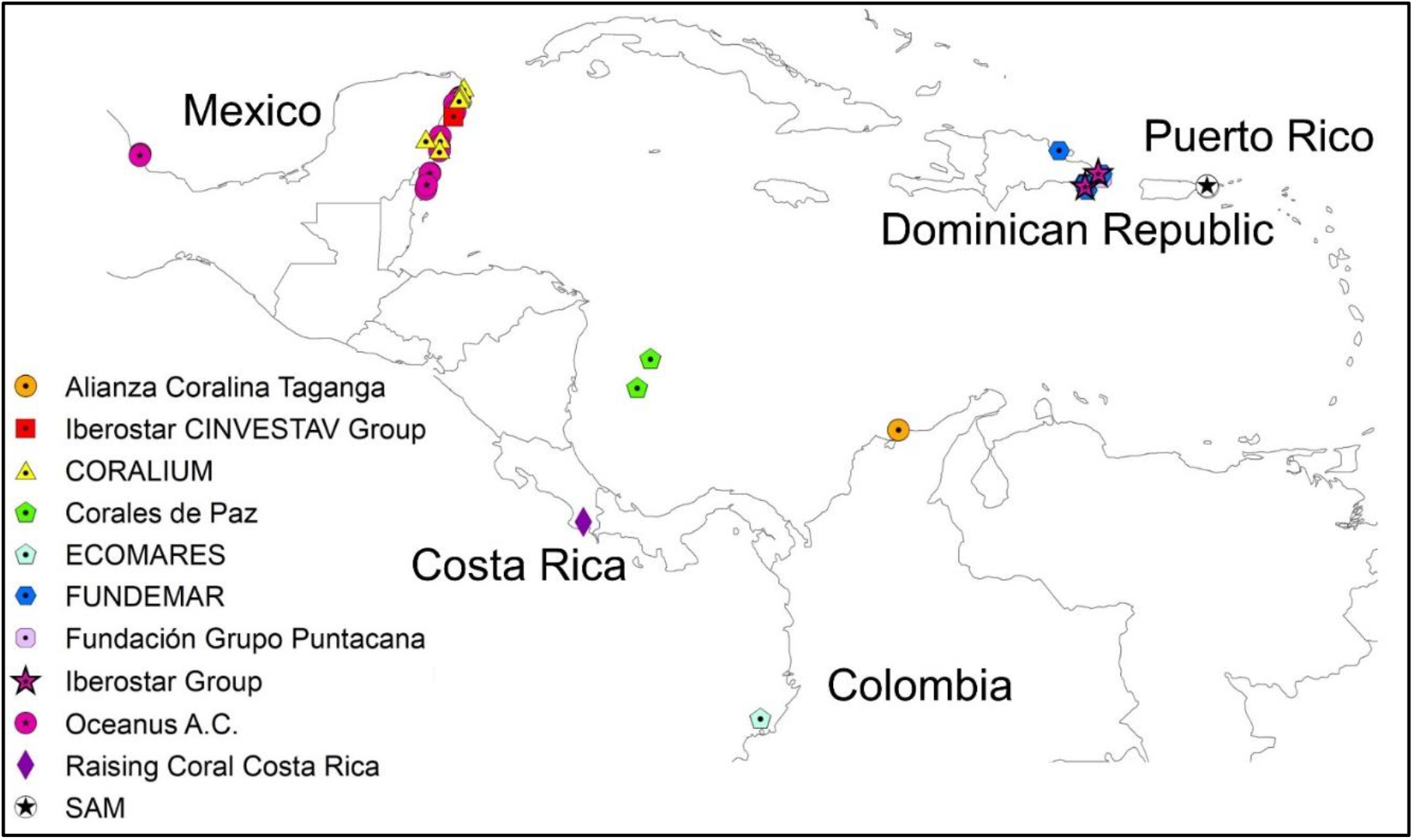
Map of coral reef restoration projects in Spanish-speaking Latin American countries and territories.

**Table 1:** Five motivation categories for carrying out coral reef restoration projects and examples.

| Motivation category | Examples |
| --- | --- |
| Biotic | Biodiversity enhancement (e.g., native species, habitat creation, ecosystem connectivity, ecological resilience) |
| Experimental | Improve restoration approaches, technologies, and methods. Answer ecologically-based research questions |
| Idealistic | Cultural reasons (e.g., recreation, tourism, medicinal/ceremonial substances, spiritual importance, aesthetic value) |
|  | Social reasons (e.g., community involvement, job creation, nature education, environmental outreach) |
|  | Political reasons (e.g., raising environmental profile) |
| Legislative | Restoration after environmental impact (e.g., ship-grounding, mining, oil spill, hurricane damage) |
|  | Biodiversity offset (e.g., threatened species, threatened ecological communities) |
| Pragmatic | Enhance ecosystem services (e.g., fisheries production) |
|  | Enhance ecosystem services (e.g., water quality improvement, pollution prevention) |
|  | Enhance ecosystem services (e.g., coastal protection, erosion control, bank stabilisation) |
|  | Enhance ecosystem services (e.g., carbon sequestration, carbon offsets) |

The objectives of coral reef restoration projects can be highly diverse and dependent on the specific project as well as its location. In this study, the restoration practitioners were asked to provide the objectives for their restoration projects, which were specific, measurable, achievable, repeatable and time-bound (SMART; [3]). We modified the six primary objectives observed by Hein et al. [39] into the following categories: 1) enhance ecosystem services for the future; 2) optimize/scale-up restoration approaches; 3) promote coral reef conservation stewardship; 4) provide alternative, sustainable livelihood opportunities; 5) reduce coral population declines and ecosystem degradation; and 6) re-establish a self-sustaining, functioning reef ecosystem.

The total estimated project cost includes both capital and operating costs. Capital costs are those used for planning, land acquisition, construction, and financing [40]. These may also include costs for laboratory/infrastructure, boats and dive equipment. Operating costs are those used for maintenance, monitoring, equipment repair and replacement [40] and may include salaries, housing for scientific/implementation teams, air for SCUBA tanks, gasoline for boat engines, and replacement of computers. Coral reef practitioners were asked to estimate the total cost for restoration interventions based on the guidelines for standardised reporting of costs for management interventions for biodiversity conservation [41] and are provided as United States Dollars (USD) per hectare of coral reef intervened per year in 2018 USD.

The project spatial extent is the coral reef area intervened by the restoration project and is reported in hectares. Spatial extent is not provided for each project since not all restoration case studies have an objective to increase the area of restored habitat. For instance, some projects are aimed at developing new restoration techniques, using coral nurseries as a tool to stimulate public awareness and engagement, for educational purposes, or as a tourist attraction.

The project duration is the time during which the restoration project has existed until the present, or the time during which the restoration cost was budgeted for and is provided in years. All projects described here are ongoing and active throughout 2019.

The feasibility is the likelihood that each specific project objective can be reached successfully with the interventions at hand and within the outlined project duration. It is ideally measured as the likelihood of success in returning the ecosystem function and resilience of an ecosystem through restoration [42]. This overall restoration project feasibility is rarely reported in the published literature because a standardised method to measure restoration success is largely missing [40]. Here, restoration practitioners estimated the feasibility of the restoration interventions they employed to achieve their specific project objectives. Feasibility is given as a ratio between 0 and 1 and can be interpreted as the likelihood of success to reach a specific objective within the duration of the restoration project. Practitioners provided a minimum, maximum and the best guess for the project feasibility.

## Results

Data from a total of 12 coral reef restoration projects carried out by practitioners in the Spanish-speaking Caribbean and Eastern Tropical Pacific were compiled and are summarised in **Table 2**. The supplementary material contains more detailed information about each restoration case study. Information was gathered from Colombia (Alianza Coralina Taganga, Corales de Paz, and ECOMARES), Costa Rica (Raising Coral Costa Rica), the Dominican Republic (FUNDEMAR, the Iberostar Group, and Fundación Grupo Puntacana), Mexico (Oceanus A.C., CORALIUM at Universidad Nacional Autónoma de México, and the Iberostar & CINVESTAV Group), and Puerto Rico (Sociedad Ambiente Marino) (**Figure 1**). Note that the Fundación Grupo Puntacana has two restoration programs of which one is focused on coral gardening (Program 1) and one is directed towards micro-fragmentation (Program 2). These were treated as independent projects for analytical purposes. The restoration projects use techniques that include direct transplantation, coral gardening, micro-fragmentation, and larval propagation (**Figure 2;** Supplementary information **Table S1**).

**Table 2:**
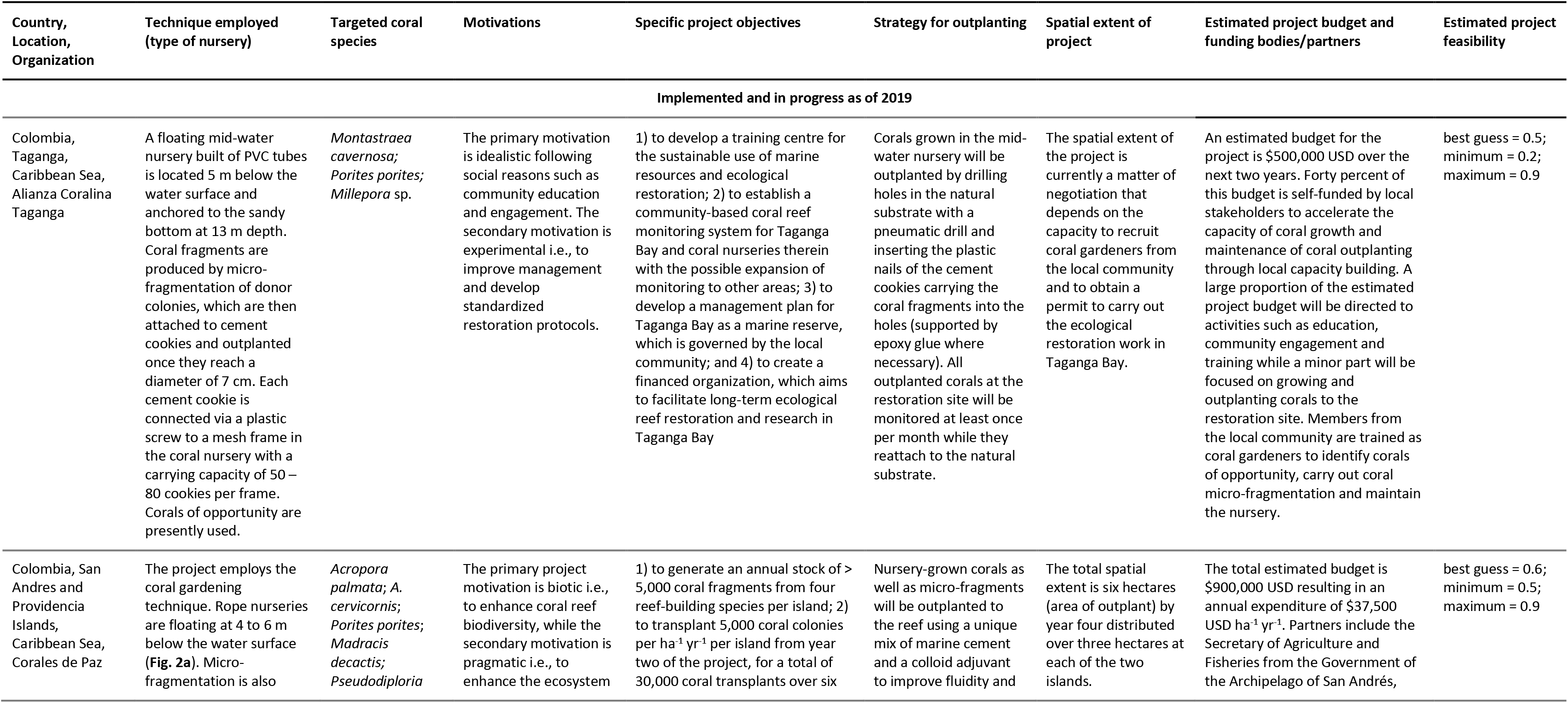
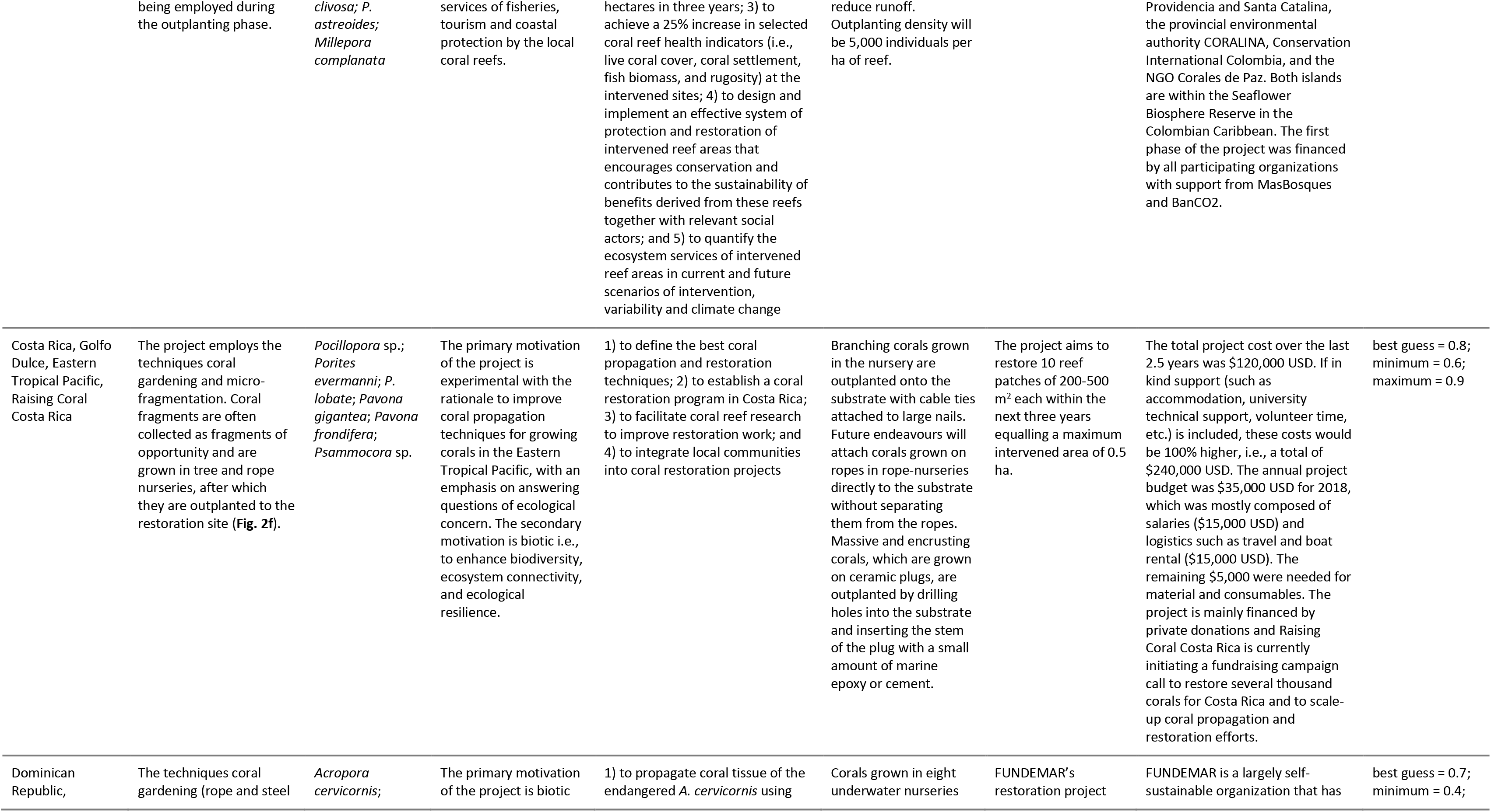
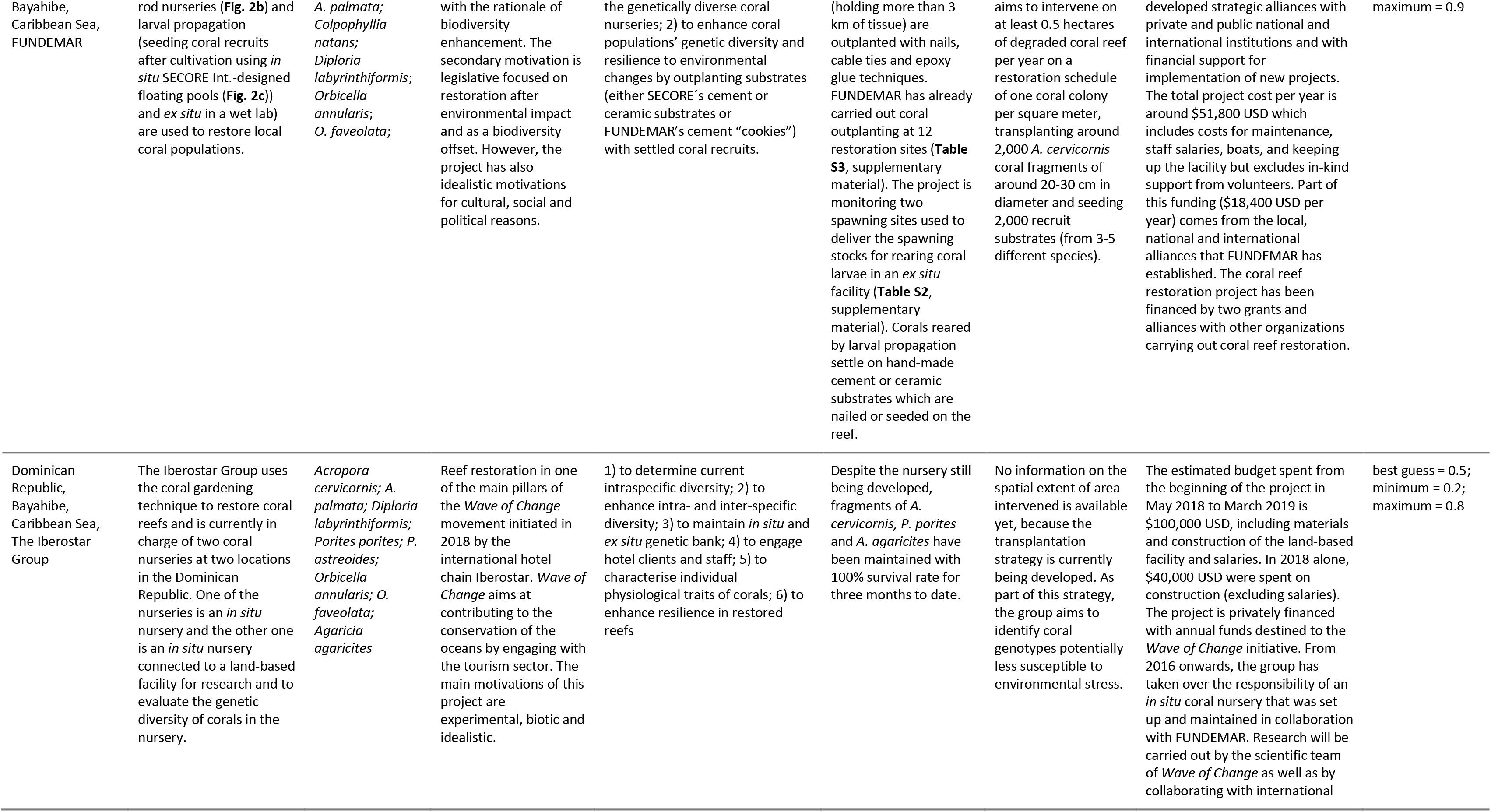
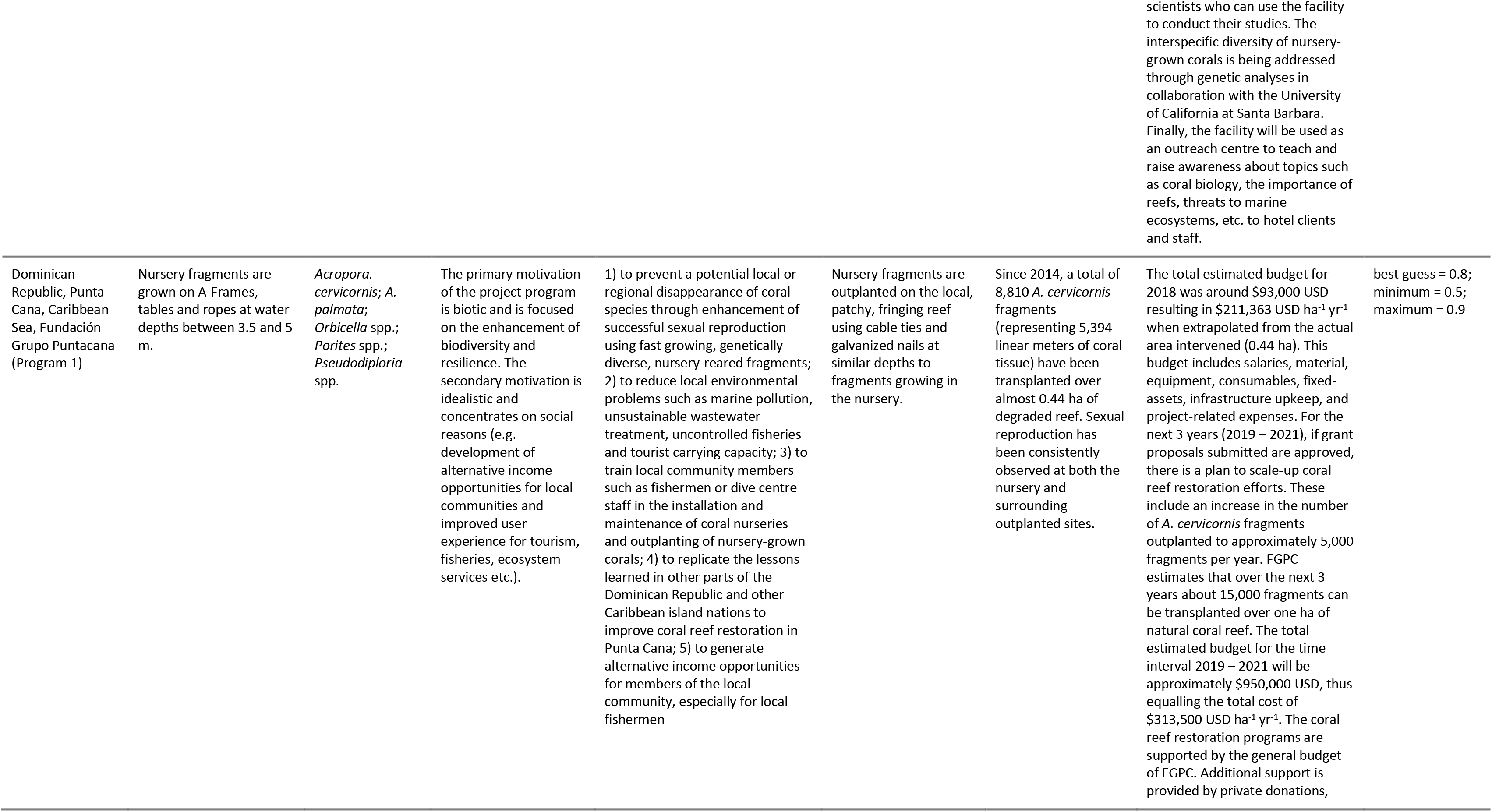
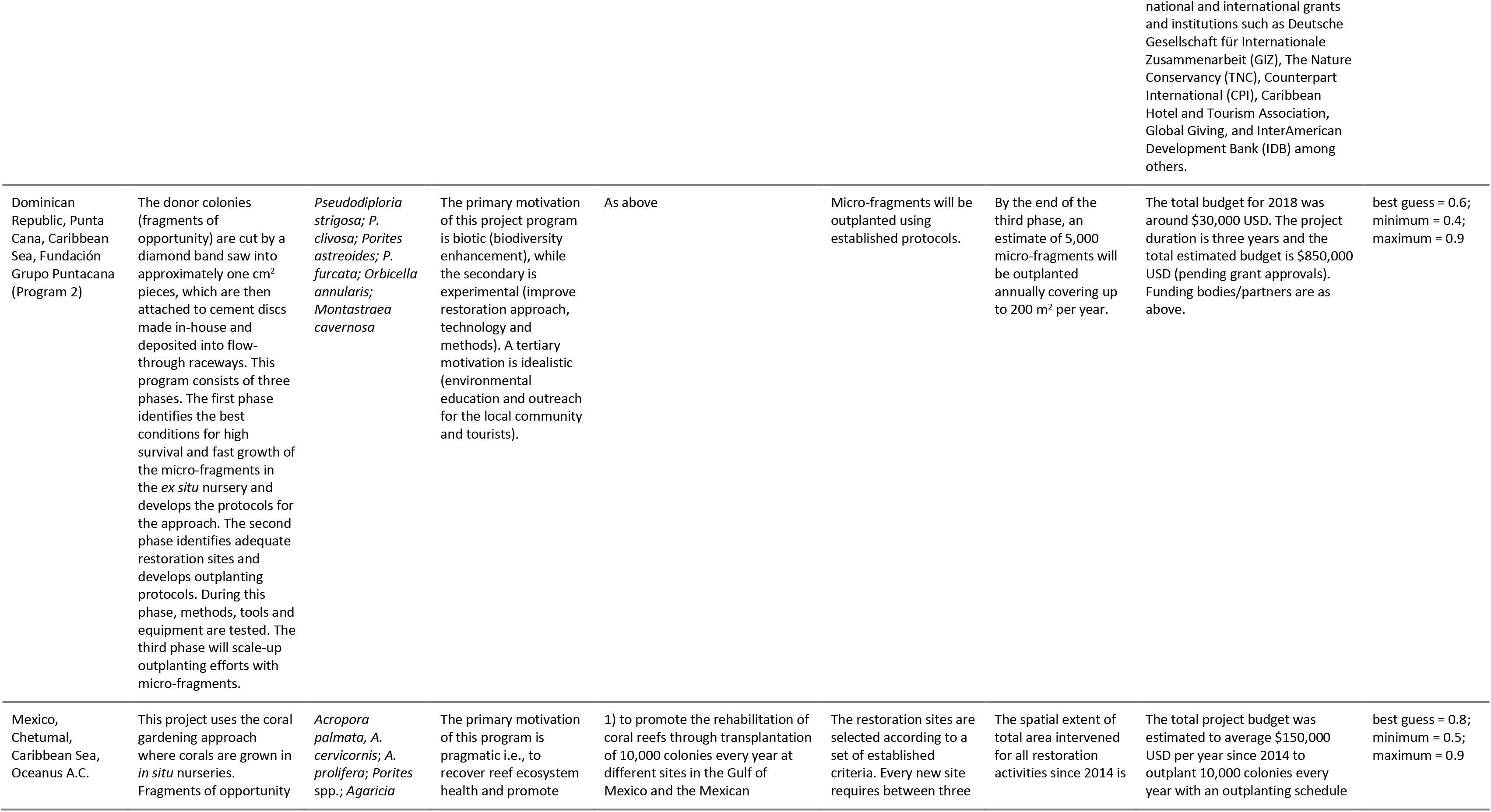
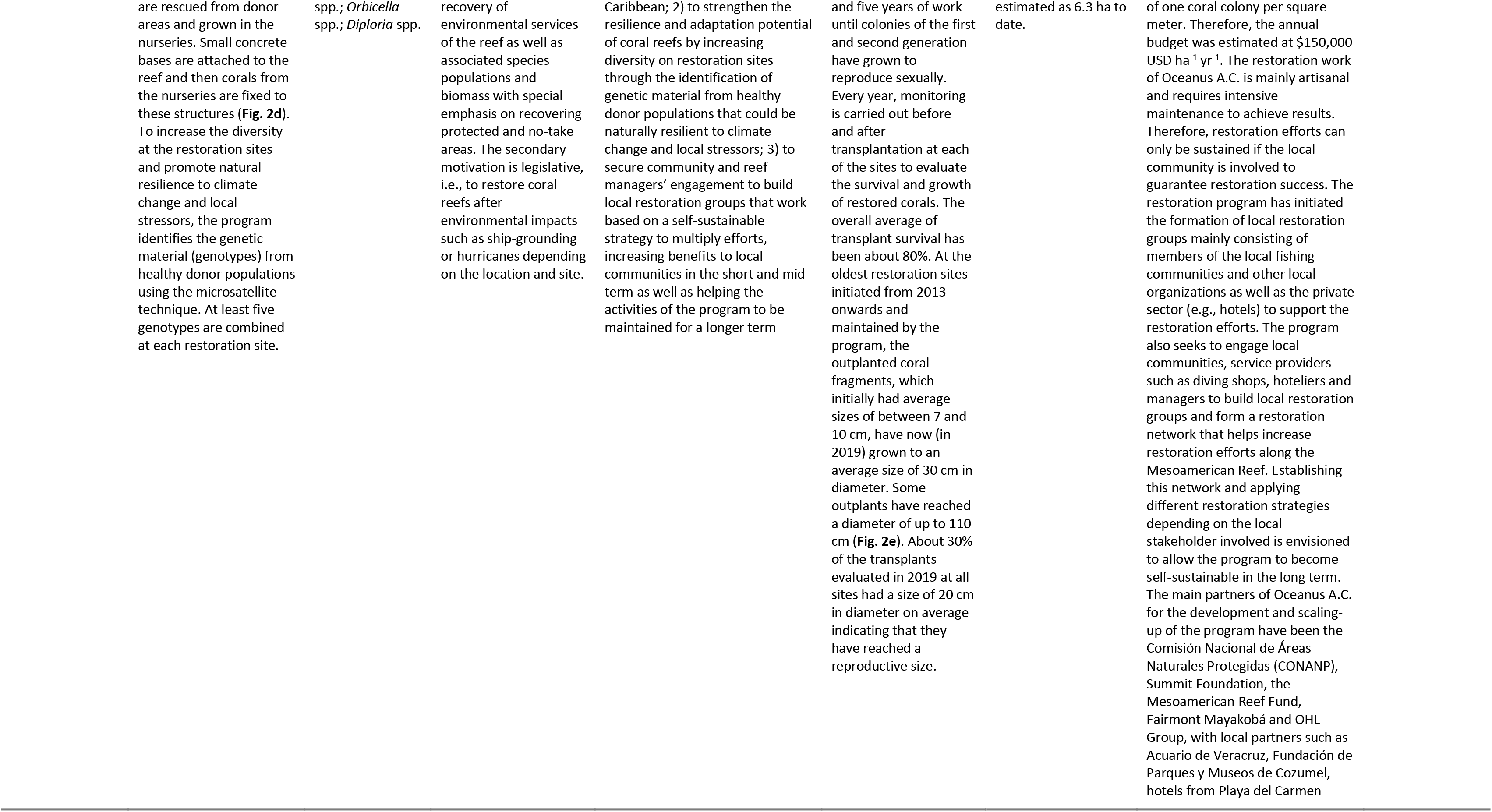
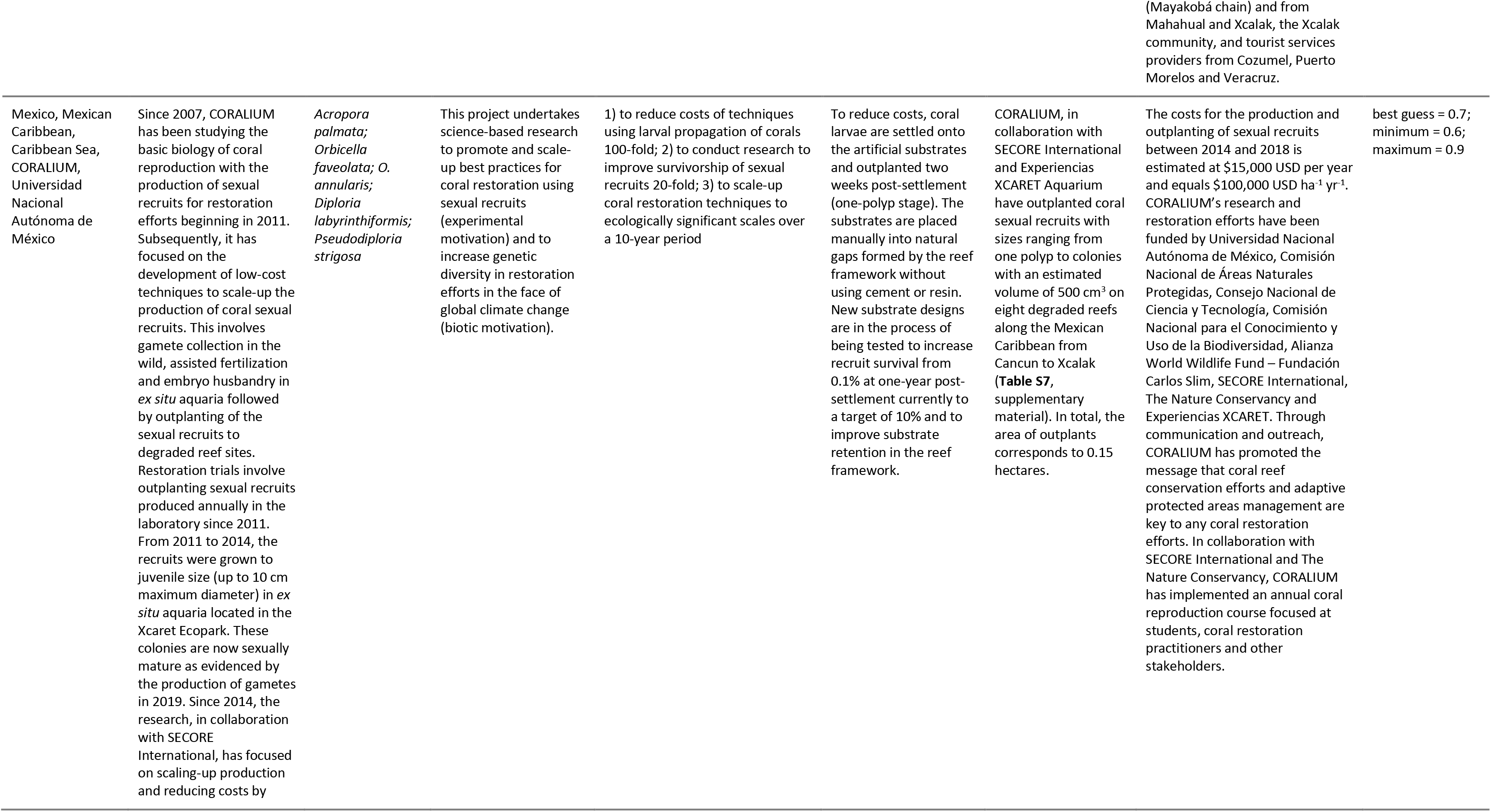
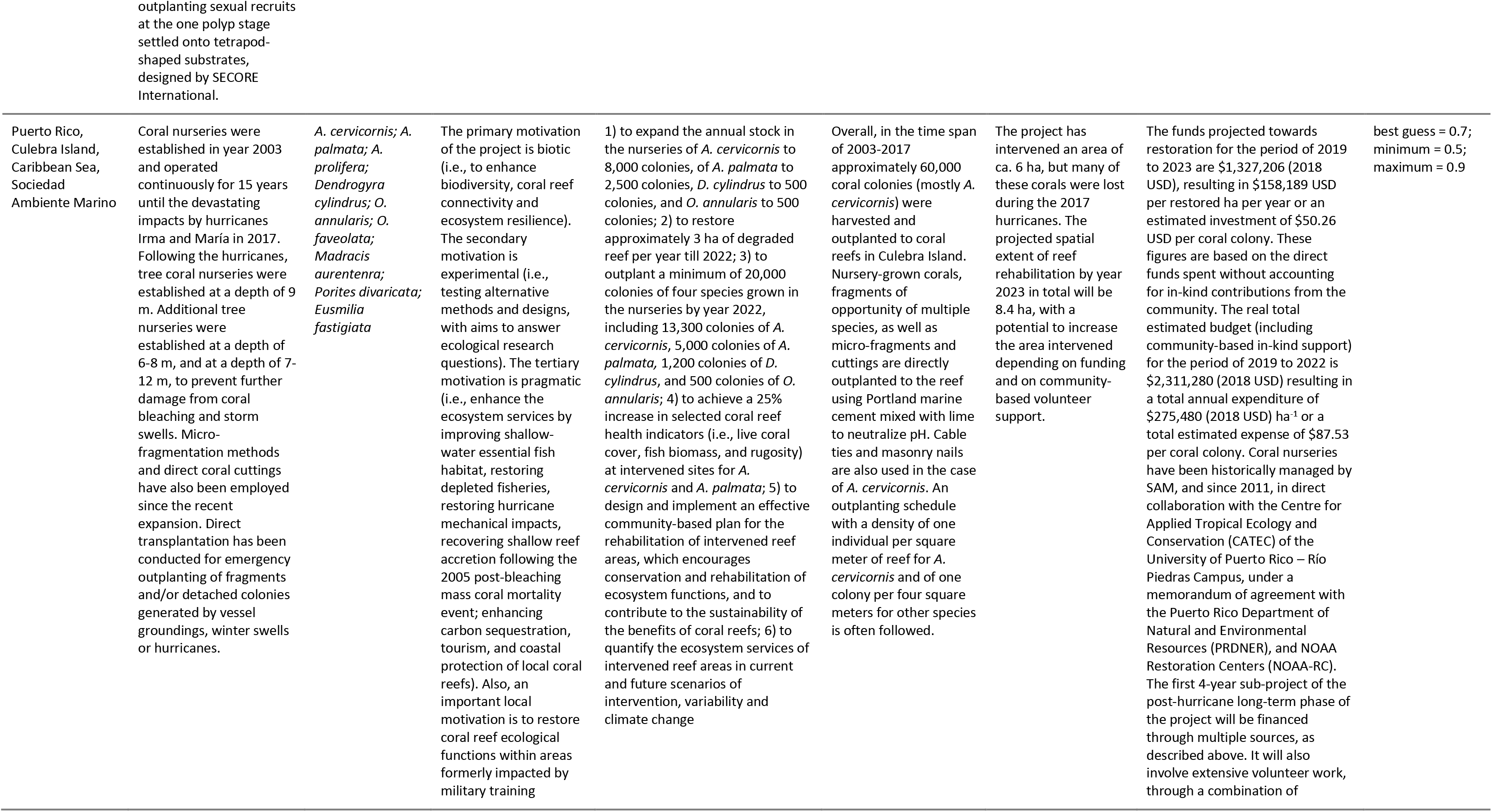
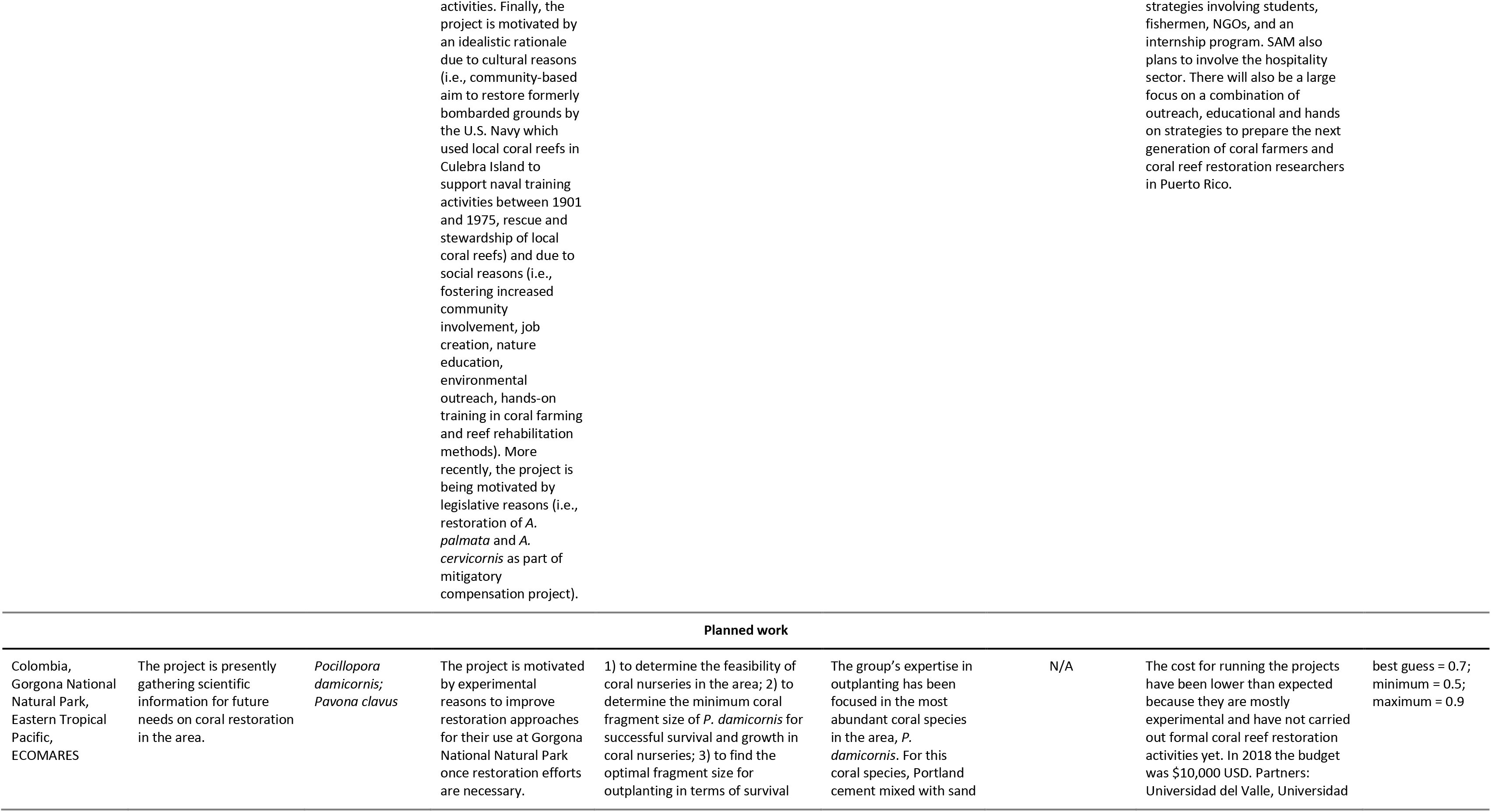
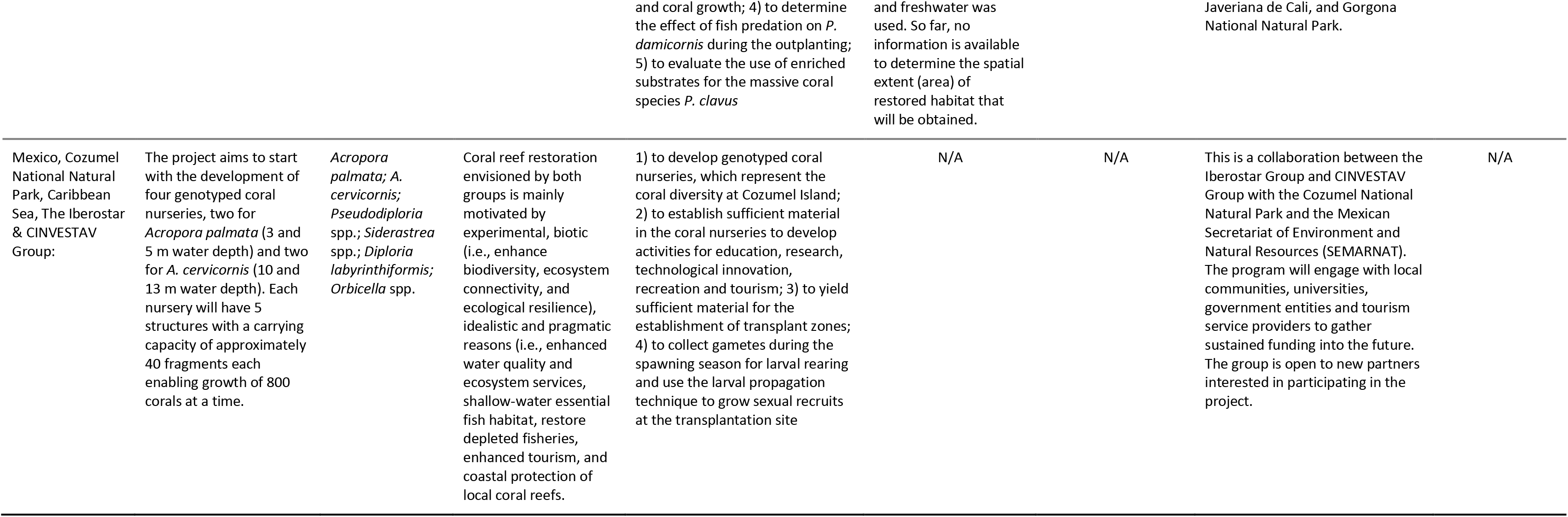
Summary of the 12 restoration projects in the Caribbean and Eastern Tropical Pacific. Cost values are given in 2018 USD. More detailed information can be found in the supplementary material. Abbreviations: Fundación Dominicana de Estudios Marinos, Inc. (FUNDEMAR), Fundación Grupo Puntacana (FGPC), and Sociedad Ambiente Marino (SAM).

**Figure 2:**
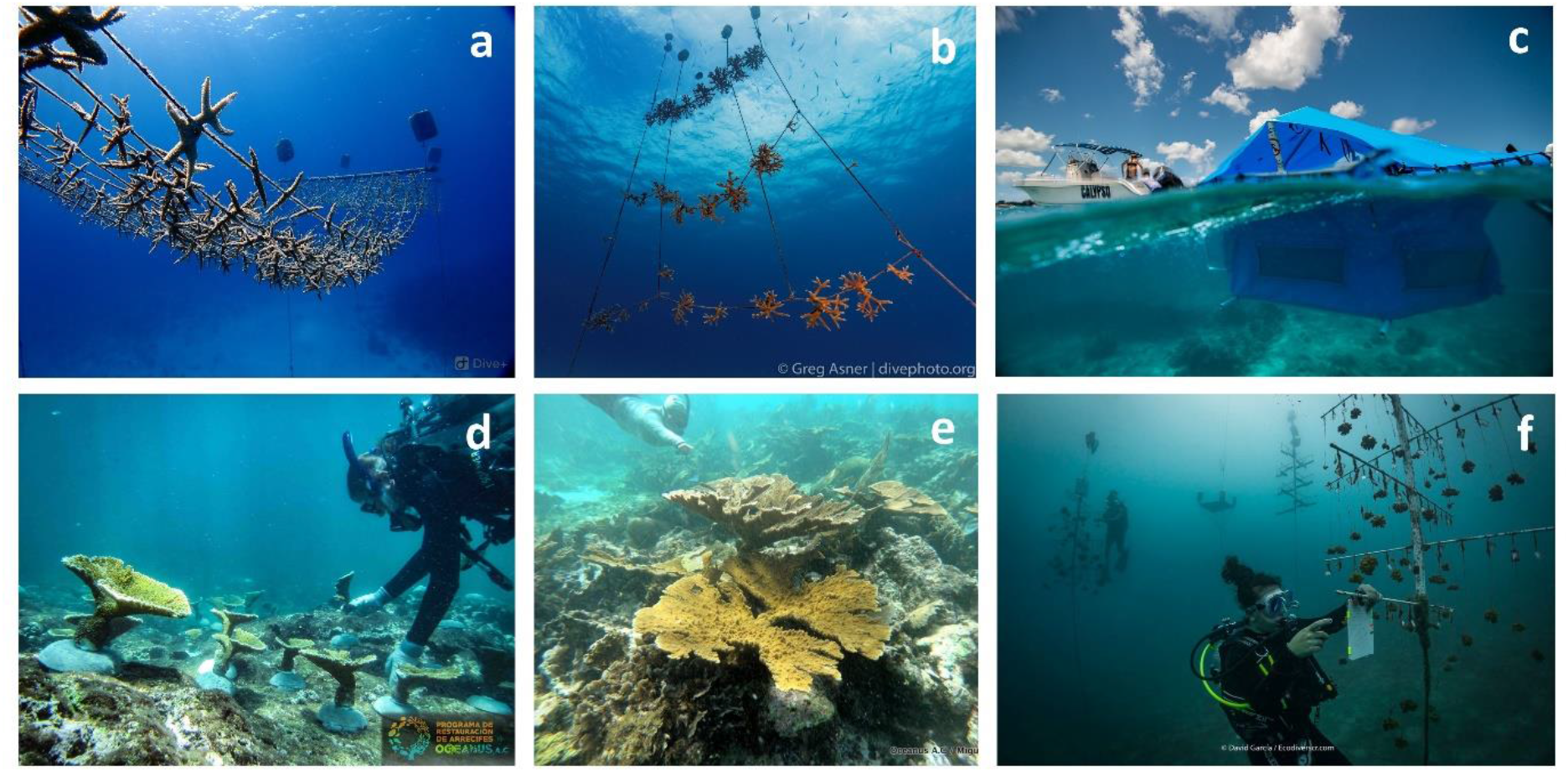
Types of nurseries described in the text. a) Floating rope nurseries used in San Andrés and Providencia islands for large-scale coral gardening (Photo: Corales de Paz); b) rope nurseries by FUNDEMAR in Dominican Republic (Photo: Greg Asner); c) FUNDEMAR’s floating *in situ* coral larvae rearing tank (Photo: Paul Selvaggio); d) Oceanus A.C. diver outplants nursery grown corals in Veracruz, Mexico (Photo: Oceanus A.C.); e) outplanted *Acropora palmata* coral in Puerto Morelos, Mexico (Photo: Oceanus A.C.); Raising Coral Costa Rica’s tree nurseries in Costa Rica (Photo: David Garcia).

The primary motivations to carry out the coral reef restoration projects are biotic and experimental to equal parts (41.7%), followed by idealistic and pragmatic reasons (both 8.3%). Biotic (36.3%) and experimental (27.3%) reasons were important secondary motivations, followed by legislative reasons (18.2%), and pragmatic/idealistic motivations (both 9.1%). All except for one of the projects reported secondary motivations. The tertiary motivations reported by 5 of the 12 projects were mainly pragmatic (80.0%) and idealistic (20.0%).

Most projects have specific objectives to optimize/scale-up restoration approaches (51.1%), followed by providing alternative, sustainable livelihood opportunities (14.9%), and then in equal parts to promote coral reef conservation stewardship and re-establish a self-sustaining, functioning reef ecosystem (12.8%). The objectives to enhance ecosystem services for the future and the reduction of population decline and ecosystem degradation accounted for only 4.2% each of the specific project objectives.

The median total cost from all projects per year is $93,000 USD (± $32,731 SE) ranging between $10,000 USD and $331,802 USD. The median spatial extent of coral reef restoration intervention is 1.0 ha (± 1.3 ha SE) ranging between 0.06 ha and 8.39 ha. Project duration was as short as 1 year and could be as long as 17 years with the median project duration of 3 years (± 1.5 years SE) to reach the project targets. Projects reported a median feasibility of 0.7 (± 0.03 SE) ranging from 0.5 to 0.8 (**Table 3**).

**Table 3:** Summary of total annual project costs, spatial extent of coral reef area intervened, project duration and feasibility from 12 case studies in the Spanish-speaking Caribbean and Eastern Tropical Pacific (Fundación Grupo Puntacana’s restoration programs were treated as two independent projects). Error is given as standard error (± SE). Abbreviation: number of observations (N).

|  | **Total cost per year**<br>**(2018 USD)** | **Spatial extent**<br>**(ha)** | **Project duration**<br>**(yrs)** | **Feasibility**<br>**(best guess)** |
| --- | --- | --- | --- | --- |
| Median | 93,000<br>(± 32,731) | 1.00<br>(± 1.30) | 3.0<br>(± 1.5) | 0.7<br>(± 0.03) |
| Min | 10,000 | 0.06 | 1 | 0.5 |
| Max | 331,802 | 8.39 | 17 | 0.8 |
| N | 11 | 7 | 12 | 11 |

## Discussion

Here we present the first comprehensive assessment of coral reef restoration projects in Spanish-speaking countries and territories of the Caribbean and Eastern Tropical Pacific (ETP), which are already being implemented or are in the initiation phase. These projects were identified through an open call for participation at the Reef Futures conference in December 2018, which aimed to bring together a large international community to develop and implement solutions to the global coral reef crisis.

We describe 12 coral reef restoration case studies in the Caribbean and Eastern Tropical Pacific that employ coral reef restoration techniques including direct transplantation, coral gardening, micro-fragmentation and larval propagation (Supplementary information, **Table S1**). With a median total project cost per year of $93,000 USD, spatial extent of 1 ha, duration of 3 years and overall project feasibility of 0.7, we show that coral reef restoration projects in these countries are more cost-effective, have overcome the barriers of scaling-up restoration interventions, are persistent through time, and have a higher likelihood of success than reported from previous literature [10, 12, 40]. For instance, the most recent published literature review on coral reef restoration presented a median value of $400,000 (2010 USD) to restore 1 ha (10,000 m^2^) of coral reef, project duration of 1 year, an area intervened of 0.01 ha (108 m^2^), and survival of restored corals as an item-based success indicator of 0.61 [10].

The objectives for coral reef restoration are often undocumented in the published literature, thus extracting data on the objectives from published papers may lead to skewed results. For example, Hein et al. [39] reviewed 83 published coral reef restoration studies and observed that 60% of the studies reported on evaluating the biological response of the coral reef ecosystem to transplantation (outplanting) as a main project objective. The remaining 40% of studies included the following objectives: 1) to accelerate reef recovery post-disturbance (18%), 2) to re-establish a self-sustaining, functioning reef ecosystem (48%), 3) to mitigate coral loss prior to a known disturbance (18%), and 4) to reduce population declines and ecosystem degradation (15%). In comparison, we observed that when data are elicited directly from restoration practitioners, most coral reef restoration projects in the Caribbean and Eastern Tropical Pacific had the following objectives: 1) to optimize or scale-up restoration approaches (51.1%), followed by 2) to provide alternative, sustainable livelihood opportunities (14.9%). Similarly, the projects presented here were mostly motivated by biotic reasons such as to enhance biodiversity and experimental reasons (both 41.7%), followed by idealistic/pragmatic reasons (both 8.3%). In contrast, most motivations to restore coral reefs extracted from the published literature were dominated by experimental reasons, such as to improve the restoration approach and answer ecological research questions (65.3%) [10]. Many restoration projects presented here focused on harnessing social or economic benefits from coral reef restoration such as involving the community through inclusion in activities or educational programs to raise awareness or to provide alternative, sustainable livelihood opportunities for local communities. An assessment of social, economic, and cultural benefits derived from the restoration of coral reefs has been largely ignored by the published literature, which has mostly concentrated on outcomes related to the ecology or described endeavours to improve restoration technology [10]. The present work is an attempt to bridge the gap between academics and practitioners. Academics tend to be more focused on small-experimental coral reef restoration attempts to answer questions of ecological concern, whereas practitioners are more focused on optimising and scaling-up restoration. Bridging the gap between academics and practitioners has been identified as critical for many fields of conservation [43, 44].

Coral reef restoration in the Caribbean and Eastern Tropical Pacific face challenges similar to those of restoration efforts elsewhere in the world. For instance, the Intergovernmental Panel on Climate Change (IPCC) concluded that, if no action is taken to reduce CO_2_ emissions, coral reefs would decline by 70-90 % with global warming of 1.5°C above pre-industrial levels, whereas virtually all coral reefs (> 99 percent) would be lost with 2°C warming within the next 50 years [45]. Thus, while actions to reduce CO_2_ emissions are drastically needed, restoration with heat resilient species is regarded as a key strategy to rehabilitate the ecological function and ecosystem services provided by coral reefs [35]. In addition to climate change, coral reef restoration in the Caribbean and ETP face other challenges such as overfishing, sedimentation, pollution, and non-sustainable coastal development [46–51]. The recent outbreak of Scleractinian Coral Tissue Loss Disease (SCTLD) has decimated coral populations and is of major concern to those attempting to restore corals in the Caribbean. Since its onset in 2017, SCTLD has caused widespread mortality of corals, especially in the Florida Reef Tract and the Gulf of Mexico [52, 53]. The vectors causing this disease or how it can be prevented are currently unknown but are most likely bacterial [52]. A further challenge to the restoration of coral reefs in the Caribbean and ETP is the apparent lack of funding and funding strategies. None of the countries have cohesive national plans for the restoration of coral reefs similar to the Reef Restoration and Adaptation Plan in Australia which has invested AUD $100 million in 2018 to develop, trial, and deploy coral reef restoration interventions for the Great Barrier Reef (GBR) [54].

Despite the impediment of limited financial resources, considerable advances in coral reef restoration, both scaling-up of interventions and optimisation of techniques, have been achieved in Colombia, Costa Rica, Dominican Republic, Mexico and Puerto Rico. For instance, one of the largest and longest running projects (18 years) has plans to restore up to 8.4 ha, requiring outplanting 10,000 corals or up to 8,000 coral settlement bases with coral larvae per year. These interventions were led by pioneering environmental NGOs and foundations, who often procured un-paid volunteers to carry out much of the work. The interventions were also enabled by strong partnerships initiated by the champion organization with universities (e.g. Universidad Nacional Autónoma de México, University of Puerto Rico, Universidad del Valle, Universidad Javeriana de Cali, Universidad de Costa Rica), conservation management bodies and regulators (e.g. Natural Parks administrations, Departments of Natural and Environmental Resources and the United States National Oceanic and Atmospheric Administration), associations (e.g. Fishers Association, Caribbean Hotel and Tourism Association), national and international business partners (e.g. SECORE International), international environmental NGOs (e.g. Conservation International, The Nature Conservancy), tourist service providers (e.g. the Iberostar Group), private donations (e.g. Global Giving), international grant schemes (e.g. from Deutsche Gesellschaft für Internationale Zusammenarbeit, Counterpart International, InterAmerican Development Bank (IDB)) and in large part with local community groups. Coral reef restoration still remains an underfunded area in the Spanish-speaking countries and territories of the Caribbean and ETP despite the ecosystem services restored coral reefs could provide for the regions such as food, tourism income, protection against storms and wave surges [55, 56], and reduction in insurance premiums by offering coastal protection [57].

There are a few caveats that need to be considered when assessing the data within the present work. First, this review does not contain an exhaustive list of interventions in the Spanish-speaking countries and territories of the Caribbean and ETP. Additional projects exist or are planned, but were not aware of, or chose to not participate in our open call. Second, the projects presented here varied in their specific objectives, best practice protocols, and monitoring, which hindered their comparison. For example, some projects were designed to improve and optimise the restoration approach (experimental projects), while others were more operational, i.e., aimed to scale-up the restoration of coral reefs by using already established restoration techniques. Furthermore, the projects used different best practice protocols or key indicators of restoration success, such as size of transplant and density of transplants which made a direct comparison between the projects difficult. Some projects lacked monitoring milestones to evaluate the survival, cover and health conditions of outplanted corals beyond year one. Yet, post-restoration monitoring is an imperative method needed to confirm that outplanted corals are self-sustaining which, from an evolutionary perspective, is the ultimate goal of any restoration effort [3–5]. Third, evaluation of the overall project feasibility or the likelihood of success to reach specific project objectives is naturally linked to local conditions and circumstances, thus may be a subjective measure directly related to the experience of the practitioner. More quantitative measures of overall project feasibility (e.g., based on measurements) would be a considerable improvement over the qualitative (derived from expert elicitation) approach.

Prior to any conservation action, a prioritisation of interventions based on decision-support frameworks is recommended to help practitioners increase their planning rigor, project accountability, stakeholder participation, transparency in decisions, and learning [58]. Cost-effectiveness analysis is such a tool that allows for the evaluation and prioritisation of conservation interventions [59]. This analysis relates the costs of a project to its key outcomes or benefits i.e., the specific measures of project effectiveness [59, 60]. Although this work includes all data required for a cost-effectiveness analysis (see Supplementary material), we considered that comparing the different projects against each other will be inappropriate given the variety of their project objectives (e.g. experimental vs. operational) and the lack of standardisation in reporting on cost, feasibility and key outcomes.

Future collaborations between academics, local communities and practitioners will be crucial if we want to achieve restoration at meaningful ecological, spatial and social scales [61]. Unfortunately, the language barrier often inhibits such collaborations. For instance, Amano et al. [62] argues that languages are still a major barrier to global science by showing that more than 35% of the knowledge in conservation is missed by those who only look at peer-reviewed literature in English. Many practitioners who carry out large-scale coral restoration projects only convey their knowledge in the form of unpublished reports and grey literature [10], which adds another level of complexity to the loss of information on restoration efforts. Here we close this gap by accessing this knowledge and overcoming the language barrier.

## Conclusions

Although not previously highlighted by the published literature, there are many coral reef restoration projects currently in progress in the Spanish-speaking countries and territories of the Caribbean and Eastern Tropical Pacific. Most of these projects are being carried out by pioneering civil organizations often in strong partnerships with universities, conservation management bodies and regulators, tourism operators, the private sector, associations, and local community groups. While coral reef restoration has been portrayed as too expensive and challenging with regards to spatial scale, duration, and success, the projects presented here have shown that many of these barriers have already been overcome. These pioneering endeavours were often possible by in-kind commitments of staff and volunteers as well as involvement of the local community and tourism operators, thus socio-economic aspects play a substantial role in coral reef restoration in the Caribbean and Eastern Tropical Pacific. Strong national plans for restoration in conjunction with national and international funding are needed to multiply the already existing activities made by Latin-American organisations to improve the health and status of coral reefs in the Caribbean and Eastern Tropical Pacific. From this compilation of data and knowledge, it is apparent that it would be beneficial for coral reef restoration practitioners in this area to coordinate their efforts with each other and make sure they are sharing and implementing their best practices protocols to standardise efforts and track restoration progress by specific, measurable, achievable and repeatable metrics of success through time.

## Acknowledgements

We would like to thank Nufar Charuvi, the pioneer driving the Alianza Coralina Taganga project, who, although no longer with us, continues to inspire by his example, dedication, and passion he served over the last decade for Colombian coral reefs and the local community. This manuscript has been developed upon in-kind time of the authors and has not received any financial support. We acknowledge the Iberostar Group for covering the open access publication fees of this manuscript.

