## Supplementary Material for "Coral reef restoration efforts in Latin American countries and territories"

**Supplementary information**

*Coral reef restoration case studies*

This overview presents 11 coral reef restoration projects carried out in Spanish-speaking countries in the Caribbean and Eastern Tropical Pacific. The projects are grouped into those that have commenced (n = 9) and interventions that were due to start in 2019 (n = 2). All projects are grouped by country, location and organization with stewardship over the project.

*Implemented projects*

Country: Colombia

Location: Taganga

Organization: Alianza Coralina Taganga

The coral reef restoration project led by Alianza Coralina Taganga is located at the Colombian Caribbean coast (11.269667° N, -74.203611° W). The project employs a floating mid-water nursery built of PVC tubes located 5 m below the water surface and which is anchored to the sandy bottom at a depth of 13 m. Coral fragments are produced by micro-fragmentation of donor colonies, which are then attached to cement cookies and outplanted once they reach a diameter of 7 cm. Each cement cookie is connected via a plastic screw to a mesh frame in the coral nursery with a carrying capacity of 50 – 80 cookies per frame. At this stage of the project, only corals of opportunity (i.e., those found detached from the substrate as a result of storms, waves or physical damage) are being used as donor colonies to reduce the impact on living corals. Members from the local community are trained as coral gardeners to identify corals of opportunity, carry out coral micro-fragmentation and maintain the nursery. The project will focus on the species *Montastraea cavernosa, Porites porites* and *Millepora* sp. The primary motivation of this project is idealistic following social reasons such as community education and engagement. The secondary motivation is experimental i.e., to improve management and develop standardized restoration protocols. This project focuses on the promotion and increase in the value of citizen participation in marine ecosystems conservation by creating activities related to coral reef restoration. An emphasis is set on improving the education of the local community and on promoting coral reef research. The specific project objectives are 1) to develop a training centre for the sustainable use of marine resources and ecological restoration; 2) to establish a community-based coral reef monitoring system for Taganga Bay and coral nurseries therein with the possible expansion of monitoring to other areas; 3) to develop a management plan for Taganga Bay as a marine reserve, which is governed by the local community; and 4) to create a financed organization, which aims to facilitate long-term ecological reef restoration and research in Taganga Bay. The project has two phases: phase one has a project duration of two years (2019 – 2020), which aims to train local community members in Taganga, while phase two relates to a long-term vision of the project leading to its institutionalisation. Here, we describe the activities of phase one. Corals grown in the mid-water nursery will be outplanted by drilling holes in the natural substrate with a pneumatic drill and inserting the plastic nails of the cement cookies carrying the coral fragments into the holes (supported by epoxy glue where necessary). All outplanted corals at the restoration site will be monitored at least once per month while they reattach to the natural substrate. The spatial extent of the project is currently a matter of negotiation that depends on the capacity to recruit coral gardeners from the local community and to obtain a permit to carry out the ecological restoration work in Taganga Bay. As a project focused on sustainability and local capability, a large proportion of the estimated project budget will be directed to activities such as education, community engagement and training while a minor part will be focused on growing and outplanting corals to the restoration site. An estimated budget for the project is $500,000 USD over the next two years. Forty percent of this budget is self-funded by local stakeholders to accelerate the capacity of coral growth and maintenance of coral outplanting through local capacity building. This budget is an estimate only; the final figure will depend on the operative capacity and will undergo constant evaluation. The best guess of estimated feasibility to achieve the four project objectives is 0.5 (minimum of 0.2 and maximum of 0.9).

Country: Colombia

Location: San Andres and Providencia Islands

Organization: Corales de Paz

In Colombia, a multi-partner collaboration was established in October 2017 to maintain two large-scale restoration projects located on San Andrés (12.494444° N, -81.735833° W) and Providencia (13.334722° N, -81.357500° W) islands. Partners include the Secretary of Agriculture and Fisheries from the Government of the Archipelago of San Andrés, Providencia and Santa Catalina, the provincial environmental authority CORALINA, Conservation International Colombia, and the NGO Corales de Paz. Both islands are within the Seaflower Biosphere Reserve in the Colombian Caribbean.

The project employs the coral gardening technique where fragments of the species *Acropora palmata*, *Acropora cervicornis*, *Porites porites*, and *Madracis decactis* are grown in rope nurseries floating at 4 to 6 meters below the water surface (**Fig. 2a**). However, micro-fragmentation is also being employed during the outplanting phase. The project has already carried out a successful pilot study to identify adequate restoration sites for the 13,468 fragments currently growing in eight nurseries (5,418 in San Andres and 8,050 in Providencia). Since October 2018, a total of 3,685 nursery-grown corals of *A. cervicornis* and *A. palmata* have been transplanted to over 2,480 square meters (outplant area; 0.248 ha) of shallow reef (<6 meters). The project has also initiated two micro-fragmentation pilot projects with *A. palmata*, *Pseudodiploria clivosa*, *Porites astreoides*, and *Millepora complanata*. The primary project motivation is biotic i.e., to enhance coral reef biodiversity, while the secondary motivation is pragmatic i.e., to enhance the ecosystem services of fisheries, tourism and coastal protection by the local coral reefs. The specific project objectives are 1) to generate an annual stock of > 5,000 coral fragments from four reef-building species per island; 2) to transplant 5,000 coral colonies per $ha^{-1}$ $yr^{-1}$ per island from year two of the project, for a total of 30,000 coral transplants over six hectares in three years; 3) to achieve a 25% increase in selected coral reef health indicators (i.e., live coral cover, coral settlement, fish biomass, and rugosity) at the intervened sites; 4) to design and implement an effective system of protection and restoration of intervened reef areas that encourages conservation and contributes to the sustainability of benefits derived from these reefs together with relevant social actors; and 5) to quantify the ecosystem services of intervened reef areas in current and future scenarios of intervention, variability and climate change. The project envisions a duration of four years of which year one was used for growing the initial coral stock in nurseries, year two was for the initial outplanting, year three is for stock maintenance, continued outplanting, and to start monitoring the intervened sites, and year four is for implementing a succession strategy. Nursery-grown corals as well as micro-fragments will be outplanted to the reef using a unique mix of marine cement and a colloid adjuvant to improve fluidity and reduce runoff [42]. Outplanting density will be 5,000 individuals per ha of reef. The total spatial extent is six hectares (area of outplant) by year four distributed over three hectares at each of the two islands. The total estimated budget is $900,000 (2018 USD) resulting in an annual expenditure of $37,500 (2018 USD) $ha^{-1}$ $yr^{-1}$. The best guess of the estimated feasibility is 0.6 (minimum of 0.5 and maximum of 0.9) based on the five specific project objectives. The first phase of the project was financed by all participating organizations with support from MasBosques and BanCO2. There is a large focus on educating, empowering and training local community members such as fishermen to maintain the coral nurseries and to continue to outplant coral fragments to the reef upon the project’s end.

Country: Costa Rica

Location: Golfo Dulce

Organization: Raising Coral Costa Rica

The civil organization Raising Coral Costa Rica is located in Golfo Dulce, on the southern Pacific coast of Costa Rica, where it maintains a coral reef restoration project (8.635544° N, -83.286635° W). The project employs the techniques coral gardening and micro-fragmentation. For the coral gardening approach, coral fragments are often collected as corals of opportunity and are grown in tree and rope nurseries, after which they are outplanted to the restoration site (**Fig. 2f**). The project focuses on the main reef building corals of the Eastern Tropical Pacific (ETP) region: *Pocillopora* sp., *Porites evermanni* and *P. lobata*, and *Pavona gigantea*. Experimental work on a smaller scale is also targeted at *Pavona frondifera* and *Psammocora* sp. The primary motivation of the project is experimental with the rationale to improve coral propagation techniques for growing corals in the ETP, with an emphasis on answering questions of ecological concern. The secondary motivation is biotic; to enhance biodiversity, ecosystem connectivity, and ecological resilience. The specific project objectives are 1) to define the best coral propagation and restoration techniques; 2) to establish a coral restoration program in Costa Rica; 3) to facilitate coral reef research to improve restoration work; and 4) to integrate local communities into coral restoration projects. Raising Coral Costa Rica has been in operation for three years (2017 - 2019) but is planned for a minimum of 10 years in total (2017 - 2026) with the possibility of an extension. Branching corals grown in the nursery are outplanted onto the substrate with cable ties attached to large nails. Future endeavours will attach corals grown on ropes in rope-nurseries directly to the substrate without separating them from the ropes. Massive and encrusting corals are outplanted by drilling holes into the substrate and inserting the stem of ceramic plugs, which carry the coral fragments with a small amount of marine epoxy or cement. The project aims to restore 10 reef patches of 200-500 $m^2$ each within the next three years equalling a maximum intervened area of 0.5 ha. The total project cost over the last 2.5 years was $120,000 USD. If in kind support (such as accommodation, university technical support, volunteer time, etc.) is included, these costs would be 100% higher, i.e., a total of $240,000 USD. The annual project budget was $35,000 USD for 2018, which was mostly composed of salaries ($15,000 USD) and logistics such as travel and boat rental ($15,000 USD). The remaining $5,000 were needed for material and consumables. The best guess of feasibility is around 0.8 (minimum 0.6 and maximum 0.9), and thus represents a high likelihood of reaching the specific project goals within the project duration. The coral species *Pocillopora* sp. has been restored with high success over the last few years. This species was nearly absent (potentially due to sedimentation) from Golfo Dulce at the onset of the project in 2017 and will be closely monitored for signs of sexual reproduction, which has so far been poorly monitored in Costa Rica. The project is mainly financed by private donations and Raising Coral Costa Rica is currently initiating a fundraising campaign call to restore several thousand corals for Costa Rica and to scale-up coral propagation and restoration efforts.

Country: Dominican Republic

Location: Bayahibe

Organization: Fundación Dominicana de Estudios Marinos, Inc. (FUNDEMAR)

The civil organization Fundación Dominicana de Estudios Marinos, Inc. (FUNDEMAR) oversees a coral reef restoration project located on the south-eastern side of the Dominican Republic (18.365881° N, -68.850397° W). The techniques coral gardening and larval propagation are used to restore local coral reefs. *Acropora cervicornis* is being restored by employing the coral gardening approach and rope nurseries (**Fig. 2b**) while *Diploria labyrinthiformis*, *A. cervicornis*, *A. palmata*, *Orbicella annularis*, *O. faveolata*, and *Colpophyllia natans* are being recovered by seeding coral recruits after cultivation. Coral larvae are reared both *in situ* using SECORE-designed floating pools (**Fig. 2c**) and *ex situ* in a wet lab. FUNDEMAR, in partnership with local and international partners, manages 8 *in situ* coral nurseries focused at the propagation of *A. cervicornis* corals in south-eastern of the Dominican Republic

(Bayahibe), and one nursery in Bavaro and another in Las Terrenas (**Table S1**, supplementary material). In the last evaluation done in 2019, FUNDEMAR hold a total of 1,873 *A. cervicornis* coral fragments in the 8 nurseries, corresponding 1,997 linear meters which had a 98% survivorship.

The primary motivation of the project is biotic with the rationale of biodiversity enhancement. The secondary motivation is legislative focused on restoration after environmental impact and as a biodiversity offset. However, the project has also idealistic motivations for cultural, social and political reasons. The project has two major objectives: 1) to propagate coral tissue of the endangered *A. cervicornis* using the genetically diverse coral nurseries: estimated feasibility 0.8 (min of 0.5, max of 1) and 2) to enhance the coral reef´s genetic diversity and resilience to environmental changes by outplanting 8,000 larval settlement bases (either SECORE´s cement or ceramic substrates or FUNDEMAR's cement "cookies") per year: estimated feasibility 0.6 (min of 0.3, max of 0.8). The project started in 2011 and is a permanent institutional program. Corals grown in the underwater nursery are outplanted by cable ties, nails and using epoxy glue where necessary. FUNDEMAR has already carried out coral outplanting at 12 restoration sites (**Table S2**, supplementary material). The project is monitoring two spawning sites used to deliver the spawning stocks for rearing coral larvae in an *ex situ* facility (**Table S3**, supplementary material). Corals reared by larval propagation either settle naturally or structures with settled coral larvae are attached by epoxy glue or nails to the substrate. FUNDEMAR's restoration project aims to intervene one hectare of degraded coral reef per year on a restoration schedule of one coral colony per square meter, transplanting around 2,000 *A. cervicornis* coral fragments of around 20-30 cm in diameter and seeding 8,000 recruit substrates (from 3-5 different species). FUNDEMAR is a largely self-sustainable organization that has developed strategic alliances with private and public national and international institutions and with financial support for implementation of new projects. The total project cost per year is around $51,800 USD which includes costs for maintenance, staff salaries, boats, and keeping up the facility but excludes in-kind support from volunteers. Part of this funding ($18,400 USD per year) comes from the local, national and international alliances that FUNDEMAR has established. The coral reef restoration project has been financed by two grants and alliances with other organizations carrying out coral reef restoration.

Country: Dominican Republic

Location: Bayahibe

Organization: The Iberostar Group

Reef restoration in one of the main pillars of the *Wave of Change* movement initiated in 2018 by the international hotel chain Iberostar. *Wave of Change* aims at contributing to the conservation of the oceans by engaging with the tourism sector. Therefore, the Iberostar Group has three overarching goals: 1) to eliminate single-use plastics in more than 120 hotels; 2) to promote sustainable fishing through acknowledging seasonal closures for breeding and reproduction of fish stocks as well as to only offer seafood from certified sustainable providers on the menus; and 3) to contribute to coastal health through conservation of seagrass meadows, mangrove forests and coral reefs. Although the goals of this movement are on a global scale, some actions are implemented differently depending on the specific location. So far, reef restoration has only been initiated in the Dominican Republic, although the Iberostar Group envisions scaling-up efforts in other locations in the Caribbean, currently in progress for Mexico in collaboration with the CINVESTAV group (see projects in Mexico). The Iberostar Group uses the coral gardening technique to restore coral reefs and is currently in charge of two coral nurseries at two locations in the Dominican Republic. One of the nurseries is an *in situ* nursery and the other one is an *in situ* nursery connected to a land-based facility for research and to evaluate the genetic diversity of corals in the nursery. The overall purpose of these two nurseries is to contribute to reef restoration practices by focusing the efforts on enhancing genetic and species diversity, and by identifying individuals that could be better suited to withstand thermal stress and hence climate change into the future. The six specific objectives of this project at the two coral reef restoration locations (Iberostar in the Bayahibe village and Iberostar Bavaro Hotel) are summarised in

**Table S4** (supplementary material). The main motivations of this project are experimental, biotic and idealistic. So far, there is no estimated duration of the restoration project, as it is still being developed under the *Wave of Change* movement. Likewise, no information on the spatial extent of area intervened is available yet, because the transplantation strategy is currently being developed. As part of this strategy, the group aims to identify coral genotypes potentially less susceptible to environmental stress.

From 2016 onwards, the group has taken over the responsibility of an *in situ* coral nursery that was set up and maintained in collaboration with the Fundación Dominicana de Estudios Marinos, Inc. (FUNDEMAR). This nursery is located in the southeast of the country, close to the Bayahibe village, and in front of the Iberostar Hotel at this location (18.339088° N, -68.826408° W). It has been placed on a sandy patch surrounded by a coral reef. The overall nursery consists of 12 structures of which three are rope nurseries, three are metal frames, one is a metal table, and five are coral nursery trees. In 2019, the *in situ* nursery produced 342 *A. cervicornis* fragments corresponding to 408 linear meters of coral tissue with 96% survivorship.

The final goal of the project is to transplant nursery-grown corals to the reef. However, because the current efforts are focused on expanding the number of coral nursery trees and enhancing intra- and interspecific diversity, so far, the restoration sites, number of transplants or project duration have not been formalised yet. The main species used in the nursery is *A. cervicornis*, although more species will be added on the longer term to address interspecific diversity. Focal species for restoration are *Diploria labyrinthiformis*, *Porites porites*, *P. astreoides*, *Orbicella annularis*, *O. faveolata*, *Agaricia agaricites*, and *A. palmata*. The interspecific diversity is being addressed through genetic analyses in collaboration with the University of California at Santa Barbara.

To support scientific restoration efforts and accomplish the specific objectives summarised in **Table S4** (supplementary material), a land-based facility is currently under construction at the Iberostar Bavaro Hotel (18.713228° N, -68.450172° W). This land-based facility will support the project by keeping a genetic bank of the coral species present in the *in situ* nursery and ensure that the unique coral genotypes are protected from storms and hurricanes, and are thus preserved into the future. The land-based facility will enable research to carry out experiments that allow for a characterization of coral individuals based on their stress tolerance to heat. Research will be carried out by the scientific team of *Wave of Change* as well as by collaborating with international scientists who can use the facility to conduct their studies. Finally, the facility will be used as an outreach centre to teach and raise awareness about topics such as coral biology, the importance of reefs, threats to marine ecosystems, etc. to hotel clients and staff. Therefore, audio-visual resources, informative signs, and entertainment activities will be implemented. Despite the nursery still being developed, fragments of *A. cervicornis, P. porites and A. agaricites* have been maintained with 100% survival rate for three months to date. All these efforts will contribute to more efficient restoration practices to guarantee higher resilience of future restored reefs. Within this project, there is also a commitment to involve the local community. For this purpose, two local fishermen have been hired and trained to help maintain the *in situ* nursery to achieve the biodiversity goals. Their actions and personal motivation to protect the ocean will make them role models for the rest of the community.

The estimated budget spent from the beginning of the project in May 2018 to March 2019 is $100,000 USD, including materials and construction of the land-based facility and salaries. In 2018 alone, $40,000 USD were spent on construction (excluding salaries). The project is privately financed with annual funds destined to the *Wave of Change* initiative. **Table S4** (supplementary material) summarises the estimated feasibility of the six specific objectives of the projects (mean best guess: 0.5 with a min of 0.2 and max of 0.8). A high feasibility is considered for the shorter-term objectives of both the land-based facility and the *in situ* nursery, due to the resources available and the recorded survival of the corals in both nurseries. Longer term objectives such as enhancing resilience of restored reefs through biodiversity approaches are considered at medium feasibility, due to the uncertainty of the factors involved in the process.

Country: Dominican Republic

Location: Punta Cana

Organization: Fundación Grupo Puntacana

The Fundación Grupo Puntacana (FGPC) was founded in 1994 to conserve and protect the natural resources of Punta Cana and to contribute to the sustainable development of the Dominican Republic. The projects and programs of the foundation are aimed at finding practical solutions to some of the local environmental issues related to marine ecosystems and sustainable development. Recently, FGPC inaugurated its Centre for Marine Innovation to lead all of its marine conservation projects including the coral reef restoration program, which hosts the largest *in situ* and the first *ex situ* coral nursery in the Dominican Republic. The nursery is located at 18.539195° N and -68.347447° W. The program applies two techniques to restore local coral reefs: since 2005 coral gardening has been used while micro-fragmentation has been implemented since 2017. The coral gardening program is mainly focused on the threatened coral *A. cervicornis* but also includes *A. palmata*. Soon, other species such as *Orbicella* spp., *Porites* spp., and *Pseudodiploria* spp. will be incorporated. Currently, over 1,500 linear meters of *A. cervicornis* tissue are being propagated at the *in situ* nursery and 12 unique genotypes are being tracked since 2011. These represent about 1,300 corals with a diameter of approximately 1.2 m.

Nursery fragments are grown on A-Frames [19], tables and ropes at water depths between 3.5 and 5 m. The primary motivation of the project program is biotic and is focused on the enhancement of biodiversity. The secondary motivation is idealistic and concentrates on social reasons (e.g. development of alternative income opportunities for local communities and improved user experience for tourism, etc.). The specific program objectives are: 1) to prevent a potential local or regional disappearance of coral species through enhancement of successful sexual reproduction using fast growing, genetically diverse, nursery-reared fragments; 2) to reduce local environmental problems such as marine pollution, unsustainable wastewater treatment, uncontrolled fisheries and tourist carrying capacity; 3) to train local community members such as fishermen or dive centre staff in the installation and maintenance of coral nurseries and outplanting of nursery-grown corals; 4) to replicate the lessons learned in other parts of the Dominican Republic and other Caribbean island nations to improve coral reef restoration in Punta Cana; and 5) to generate alternative income opportunities for members of the local community, especially for local fishermen. The Fundación Grupo Puntacana has two programs in place, one of which uses the coral gardening technique and the other one employs the micro-fragmentation approach.

Program 1: Nursery fragments are outplanted on the local, patchy, fringing reef using cable ties and galvanized nails at similar depths to fragments growing in the nursery. Since 2014, a total of 8,810 *A. cervicornis* fragments (representing 5,394 linear meters of coral tissue) have been transplanted over almost 0.44 ha of degraded reef. Sexual reproduction has been consistently observed at both the nursery and surrounding outplanted sites. The total estimated budget for 2018 was around $93,000 USD resulting in $211,363 USD $ha^{-1}$ $yr^{-1}$ when extrapolated from the actual area intervened (0.44 ha). This budget includes salaries, material, equipment, consumables, fixed-assets, infrastructure upkeep, and project-related expenses. For the next 3 years (2019 – 2021), if grant proposals submitted are approved, there is a plan to scale-up coral reef restoration efforts. These include an increase in the number of *A. cervicornis* fragments outplanted to approximately 5,000 fragments per year. FGPC estimates that over the next 3 years about 15,000 fragments can be transplanted over one ha of natural coral reef. The total estimated budget for the time interval 2019 – 2021 will be approximately $950,000 USD, thus equalling the total cost of $313,500 USD $ha^{-1}$ $yr^{-1}$. The best guess of the estimated feasibility for reaching the five project objectives is 0.8 (minimum of 0.5 and maximum 0.9), if grant proposals submitted are approved.

Program 2: The micro-fragmentation program is currently focused on the species *Pseudodiploria strigosa*, *P. clivosa*, *Porites astreoides*, and *P. furcata*. However, the project envisions including *Orbicella annularis* and *Montastraea cavernosa* as well as a couple of other species. The donor colonies (fragments of opportunity) are cut by a diamond band saw into approximately one $cm^2$

pieces, which are then attached to cement discs made in-house and deposited into flow-through raceways. This program consists of three phases. The first phase identified the best conditions for high survival and fast growth of the micro-fragments in the *ex situ* nursery and developed the protocols for the approach. This phase is complete. The second phase, beginning in 2019, will identify adequate restoration sites and develop outplanting protocols. During this second phase, methods, tools and equipment will be tested. The third phase will scale-up outplanting efforts with micro-fragments. By the end of the third phase, an estimate of 5,000 micro-fragments will be outplanted annually using established protocols, covering up to 200 $m^2$ per year. The primary motivation of this project program is biotic (biodiversity enhancement), while the secondary is experimental (improve restoration approach, technology and methods). A tertiary motivation is idealistic (environmental education and outreach for the local community and tourists). The total budget for 2018 was around $30,000 USD. The project duration is three years and the total estimated budget is $850,000 USD (pending grant approvals). The best guess of the estimated feasibility for program 2 is 0.6 (minimum of 0.4 and maximum 0.9). Both programs have the same specific objectives.

The coral reef restoration programs are supported by the general budget of Fundación Grupo Puntacana. Additional support is provided by private donations, national and international grants and institutions such as Deutsche Gesellschaft für Internationale Zusammenarbeit (GIZ), The Nature Conservancy (TNC), Counterpart International (CPI), Caribbean Hotel and Tourism Association, Global Giving, and InterAmerican Development Bank (IDB) among others.

Country: Mexico

Location: Chetumal

Organization: Oceanus A.C.

Oceanus A.C. is a Mexican non-governmental organization (non-profit) based in Chetumal, in the state of Quintana Roo, that develops projects for coral reef conservation. Its mission is to develop and apply techniques that enhance coral reef resilience, implement conservation interventions and promote the sustainable use of coastal and marine resources. Together with its partners, Oceanus A.C. has designed and implemented a Coral Reef Restoration Program for the reefs of the Gulf of Mexico and the Mexican Caribbean. This program focuses on increasing the adaptation and recovery potential of coral reefs living at the reef crest through the establishment of rehabilitation sites. This project uses the coral gardening approach where corals are grown in *in situ* nurseries. The primary motivation of this program is pragmatic i.e., to recover reef ecosystem health and promote recovery of environmental services of the reef as well as associated species populations and biomass with special emphasis on recovering protected and no-take areas. The secondary motivation is legislative, i.e., to restore coral reefs after environmental impacts such as ship-grounding or hurricanes depending on the location and site.

Between 2009 and 2018, the activities of Oceanus A.C. have focused on the rehabilitation of sites with *Acropora palmata* as a key reef building species. The habitat targeted was the reef crest because it dissipates 86% of wave energy and has an important role in coastal protection [43]. In 2019, the program plans to extend its efforts to other habitats such as the back reef and coastal reefs and to add other species of *Acropora* spp. and other genera according to their preferred environment (*A. cervicornis*, *A. prolifera*, *Porites* spp., *Agaricia* spp., *Orbicella* spp., and *Diploria* spp.). The main restoration techniques include coral gardening, which involves the construction and installation of coral nurseries for stabilization of fragments of opportunity rescued from donor areas. Thousands of colonies grown in these nurseries are being outplanted to the reef each year by first attaching small concrete bases to the reef and then fixing corals from the nurseries to these structures (**Fig. 2d**). To increase the diversity at the restoration sites and promote natural resilience to climate change and local stressors, the program identifies the genetic material (genotypes) from healthy donor populations using the microsatellite technique [44]. At least five genotypes are combined at each restoration site. The program also seeks to engage local communities, service providers such as diving shops, hoteliers and managers to build local restoration groups and form a restoration network that helps increase restoration efforts along the Mesoamerican Reef. Establishing this network and applying different restoration strategies depending on the local stakeholder involved is envisioned to allow the program to become self-sustainable in the long term.

The program has three specific objectives: 1) to promote the rehabilitation of coral reefs through transplantation of 10,000 colonies every year at different sites in the Gulf of Mexico and the Mexican Caribbean; 2) to strengthen the resilience and adaptation potential of coral reefs by increasing diversity on restoration sites through the identification of genetic material from healthy donor populations that could be naturally resilient to climate change and local stressors; and 3) to secure community and reef managers' engagement to build local restoration groups that work based on a self-sustainable strategy to multiply efforts, increasing benefits to local communities in the short and mid-term as well as helping the activities of the program to be maintained for a longer term. Currently (2018-2019), several locations for coral reef restoration are being developed by the program: at Veracruz, Xcalak, Playa del Carmen, Puerto Morelos, Sian Ka'an and just recently at Cozumel (see **Table S5**, supplementary material for all restoration sites). The restoration sites are selected according to a set of established criteria. Every new site requires between three and five years of work until colonies of the first and second generation have grown to reproduce sexually. Every year, monitoring is carried out before and after transplantation at each of the sites to evaluate the survival and growth of restored corals. The overall average of transplant survival has been about 80%. At the oldest restoration sites initiated from 2013 onwards and maintained by the program, the outplanted coral fragments, which initially had average sizes of between 7 and 10 cm, have now (in 2019) grown to an average size of 30 cm in diameter. Some outplants have reached a diameter of up to 110 cm (**Fig. 2e**). About 30% of the transplants evaluated in 2019 at all sites had a size of 20 cm in diameter on average indicating that they have reached a reproductive size [45].

The restoration work of Oceanus A.C. is mainly artisanal and requires intensive maintenance to achieve results. Therefore, restoration efforts can only be sustained if the local community is involved to guarantee restoration success. The restoration program has initiated the formation of local restoration groups mainly consisting of members of the local fishing communities and other local organizations as well as the private sector (e.g., hotels) to support the restoration efforts. The formation of these groups also allows for increasing the local capacities by involving volunteers and visitors in the restoration activities. Therefore, Oceanus A.C. has designed a certification for ‘Restoration Program Guides’. To be certified, interested participants must formally register and commit to the restoration project. The vision is to maintain the restoration activities at those sites while having the support of local groups who increase the awareness of visitors and volunteers on the importance of the program for local coral reefs. Within these activities, options for funding are sought for every local restoration group to achieve self-sustaining restoration efforts at each site over the long term. The main partners of Oceanus A.C. for the development and scaling-up of the program have been the Comisión Nacional de Áreas Naturales Protegidas (CONANP), Summit Foundation, the Mesoamerican Reef Fund, Fairmont Mayakobá and OHL Group, with local partners such as Acuario de Veracruz, Fundación de Parques y Museos de Cozumel, hotels from Playa del Carmen (Mayakobá chain) and from Mahahual and Xcalak, the Xcalak community, and tourist services providers from Cozumel, Puerto Morelos and Veracruz.

The total project budget was estimated to average $150,000 USD per year since 2014 to outplant 10,000 colonies every year with an outplanting schedule of one coral colony per square meter. Therefore, the annual budget was estimated at $150,000 USD $ha^{-1}$ $yr^{-1}$. The spatial extent of total area intervened for all restoration activities since 2014 is estimated as 6.3 ha to date. The best guess of feasibility to reach the three project objectives based on the project’s experiences with local stakeholders is around 0.8 (minimum 0.5 and maximum 0.9).

Country: Mexico

Location: Mexican Caribbean

Organization: Universidad Nacional Autónoma de México, Integrative Reef Conservation Research Laboratory (CORALIUM)

The Integrative Reef Conservation Research Lab (CORALIUM), based at the Universidad Nacional Autónoma de Mexico (UNAM) campus in Puerto Morelos, Mexico, undertakes science-based research to promote and scale-up best practices for coral restoration using sexual recruits (experimental motivation) and to increase genetic diversity in restoration efforts in the face of global climate change (biotic motivation). Specifically, CORALIUM's objectives are 1) to reduce costs of techniques using larval propagation of corals 100-fold; 2) to conduct research to improve survivorship of sexual recruits 20-fold; and 3) to scale-up coral restoration techniques to ecologically significant scales over a 10-year period. To achieve these objectives, CORALIUM collaborates with national and international researchers, NGOs, businesses and works closely with the Mexican Commission for Natural Protected Areas.

Since 2007, CORALIUM has been studying the basic biology of coral reproduction with the production of sexual recruits for restoration efforts beginning in 2011. Subsequently, it has focused on the development of low-cost techniques to scale-up the production of coral sexual recruits. This involves gamete collection in the wild, assisted fertilization and embryo husbandry in *ex situ* aquaria followed by outplanting of the sexual recruits to degraded reef sites. Restoration trials involve outplanting sexual recruits produced annually in the laboratory since 2011. From 2011 to 2014, the recruits were grown to juvenile size (up to 10 cm maximum diameter) in *ex situ* aquaria located in the Xcaret Ecopark. These colonies are now sexually mature as evidenced by the production of gametes in 2019. Since 2014, the research, in collaboration with SECORE International, has focused on scaling-up production and reducing costs by outplanting sexual recruits at the one polyp stage settled onto tetrapod-shaped substrates, designed by SECORE International. Although the production and outplanting of sexual recruits of *Acropora palmata* have been the focus, CORALIUM has also successfully worked with other important reef-building corals including *Orbicella faveolata, O. annularis*, *Diploria labyrinthiformis*, and *Pseudodiploria strigosa.*

CORALIUM, in collaboration with SECORE International and Experiencias XCARET Aquarium have outplanted coral sexual recruits with sizes ranging from one polyp to colonies with an estimated volume of 500 $cm^3$ on eight degraded reefs along the Mexican Caribbean from Cancun to Xcalak (**Table S6**, supplementary material). In total, the area of outplants corresponds to 0.15 hectares. To reduce costs, coral larvae are settled onto the artificial substrates and outplanted two weeks post-settlement (one-polyp stage). The substrates are placed manually into natural gaps formed by the reef framework without using cement or resin. New substrate designs are in the process of being tested to increase recruit survival from 0.1% at one-year post-settlement currently to a target of 10% and to improve substrate retention in the reef framework. The costs for the production and outplanting of sexual recruits between 2014 and 2018 is estimated at $15,000 USD per year and equals $100,000 USD $ha^{-1}$ $yr^{-1}$. The best guess of feasibility for reaching CORALIUM's objectives within 10 years is around 0.7 (minimum 0.6 and maximum 0.9). CORALIUM's research and restoration efforts have been funded by Universidad Nacional Autónoma de México, Comisión Nacional de Áreas Naturales Protegidas, Consejo Nacional de Ciencia y Tecnología, Comisión Nacional para el Conocimiento y Uso de la Biodiversidad, Alianza World Wildlife Fund – Fundación Carlos Slim, SECORE International, The Nature Conservancy and Experiencias XCARET.

Through communication and outreach, CORALIUM has promoted the message that coral reef conservation efforts and adaptive protected areas management are key to any coral restoration efforts. In collaboration with SECORE International and The Nature Conservancy, CORALIUM has implemented an annual coral reproduction course focused at students, coral restoration practitioners and other stakeholders. One of the most important results of this capacity-building practice has been the creation of a network throughout the Caribbean to scale-up coral restoration on a wider level: this includes Expedition Akumal (Akumal, Mexico), Cozumel Coral Reef Restoration (Cozumel, Mexico),

Mexican Fisheries Department (Puerto Morelos, Mexico), the University of Belize (Calabash Caye, Belize), FUNDEMAR (Punta Cana, Dominican Republic) and the National Aquarium of Cuba (Cuba) who have included larval propagation techniques into their restoration programs. In the near future, programs are also expected to be developed in Colombia and Costa Rica. Additionally, CORALIUM has produced educational material such as technical manuals [31, 46], and informational guides and pamphlets about the reproductive biology of corals and coral restoration.

Country: Puerto Rico

Location: Culebra Island

Organization: Sociedad Ambiente Marino, Community-Based Coral Aquaculture and Reef Rehabilitation Program - Hope for the Reef

In 2003, the non-governmental organization Sociedad Ambiente Marino (SAM), in collaboration with the Culebra Island Fishers Association, Coralations, Caborrojeños Pro Salud y Ambiente, and the University of Puerto Rico (UPR), launched the Community-Based Coral Aquaculture and Reef Rehabilitation Project in Culebra Island. The island community is located 27 km off northeast Puerto Rico, in the northeast of the Caribbean Sea. The program has undergone major adaptations as a result of impacts related to adverse environmental factors. These include localized runoff impacts, sea surface warming and mass coral bleaching (2005), and multiple category five hurricanes. For instance, hurricanes Irma and María in September 2017 nearly wiped out the entire project and eliminated over 60,000 outplanted colonies [47]. This major impact of environmental factors led to the implementation of adaptive management and change in the maintenance of coral nurseries. In the beginning, three original coral nursery sites were located at Bahía Tamarindo (BTA) (18.315272° N, -65.318129° W), Punta Melones (PME) (18.305053° N, -65.312645° W) and Punta Soldado (PSO) (18.280184° N, -65.287553° W). The original nurseries consisted of plastic-covered wire mesh cages, which were followed by line nurseries. Line nurseries were more successful for the restoration sites [48]. A total of 15 different methods have been tested through the project, including a wide variety of benthic (i.e., plastic-covered wire mesh arrays, “A” frames, PVC frames, PVC habitat structures, wire mesh cylinders) and floating coral nurseries (i.e., table-line nurseries, and tree nurseries). The sites BTA, PME, and PTC are located within the Canal Luis Peña no-take Natural Reserve. PSO is open to fishing. The original nurseries were located over sandy or rubble bottom at a water depth of 3-5 m. The PME site was abandoned in 2005 following a massive coral bleaching event. Coral bleaching in combination with increased runoff from adjacent construction sites along the coast resulted in major coral mortality at this site. Following the massive coral bleaching event, the coral nurseries at PSO were relocated to a depth of 6-8 m. Coral nurseries were re-established at PME in 2011 but were eliminated again in 2013 after storm swells and runoff significantly impacted the location in 2011 and 2012 [48]. Following impacts by hurricanes Irma and María in 2017, tree coral nurseries at BTA were established at a depth of 9 m. Additional tree nurseries were established at Punta Tamarindo Chico (PTC) (18.310547° N, -65.317597° W) at a depth of 6-8 m, and at PSO at a depth of 7-12 m, to prevent further damage from coral bleaching and storm swells. At this stage and after post-hurricane reconstruction of coral nurseries, the project was renamed *Hope for the Reef* (‘Esperanza para el Arrecife’). Coral nurseries have been historically managed by SAM, and since 2011, in direct collaboration with the Centre for Applied Tropical Ecology and Conservation (CATEC) of the University of Puerto Rico – Río Piedras Campus, under a memorandum of agreement with the Puerto Rico Department of Natural and Environmental Resources (PRDNER), and NOAA Restoration Centre (NOAA-RC).

The project mostly employs the technique coral gardening, where fragments of the species *A. cervicornis* and *A. palmata* are grown in tree nurseries to date. Floating units have been located at a water depth between 4 to 10 m. Recent major hurricane impacts led to the generation of multiple fragments of opportunity. Additional species are now grown in the nurseries including *Dendrogyra cylindrus*, *O. annularis*, *O. faveolata*, *Madracis aurentenra*, *Porites divaricata*, and *Eusmilia fastigiata*.

Micro-fragmentation methods and direct coral cuttings have also been employed since the recent expansion. Direct transplantation has been conducted for emergency outplanting of fragments and/or detached colonies generated by vessel groundings, winter swells or hurricanes. Overall, in the time span of 2003-2017 approximately 60,000 coral colonies (mostly *A. cervicornis*) were harvested and outplanted to coral reefs in Culebra Island. The project has intervened an area of ca. 6 ha.

The primary motivation of the project is biotic (i.e., to enhance biodiversity, coral reef connectivity and ecosystem resilience). The secondary motivation is experimental (i.e., testing alternative methods and designs, with aims to answering ecological research questions). The tertiary motivation is pragmatic (i.e., enhance the ecosystem services by improving shallow-water essential fish habitat, restoring depleted fisheries, enhancing carbon sequestration, tourism, and coastal protection of local coral reefs). Also, an important local motivation is to restore coral reef ecological functions within areas formerly impacted by military training activities [47, 48]. Finally, the project is motivated by an idealistic rationale due to cultural reasons (i.e., community-based aim to restore formerly bombarded grounds by the U.S. Navy which used local coral reefs in Culebra Island to support naval training activities between 1901 and 1975, rescue and stewardship of local coral reefs) and due to social reasons (i.e., fostering increased community involvement, job creation, nature education, environmental outreach, hands-on training in coral farming and reef rehabilitation methods). More recently, the project is being motivated by legislative reasons (i.e., restoration of *A. palmata* and *A. cervicornis* as part of mitigatory compensation project).

After the impacts on the restoration sites caused by category five hurricanes Irma and María in 2017, the original project objectives were modified to the following: 1) to expand the annual stock in the nurseries of *A. cervicornis* to 8,000 colonies, of *A. palmata* to 2,500 colonies, *D. cylindrus* to 500 colonies, and *O. annularis* to 500 colonies; 2) to restore approximately 3 ha of degraded reef per year till 2022; 3) to outplant a minimum total of 20,000 colonies of four species grown in the nurseries by year 2022, including 13,300 colonies of *A. cervicornis*, 5,000 colonies of *A. palmata*, 1,200 colonies of *D. cylindrus*, and 500 colonies of *O. annularis*; 4) to achieve a 25% increase in selected coral reef health indicators (i.e., live coral cover, fish biomass, and rugosity) at intervened sites for *A. cervicornis* and *A. palmata*; 5) to design and implement an effective community-based plan for the rehabilitation of intervened reef areas, which encourages conservation and rehabilitation of ecosystem functions, and to contribute to the sustainability of the benefits of coral reefs; and 6) to quantify the ecosystem services of intervened reef areas in current and future scenarios of intervention, variability and climate change. This will be achieved by combining traditional *in situ* low-tech coral gardening, but also by incorporating micro-fragmenting and cuttings of outplanted corals in order to increase the number of outplanted colonies. In addition, the production of additional peer reviewed publications is also a paramount goal for SAM which goes beyond this project.

The post-hurricanes project phase envisions a duration of ~20 years, divided into sub-projects of four years each, of which year one is used for growing the initial coral stock in nurseries, year two is for stock maintenance and for the initial outplanting, year three is for stock maintenance, continued outplanting to additional locations, and to start monitoring at the intervened sites, and year four is for implementing the sustainability strategy, renewing coral nursery structures, rotating locations, micro-fragmenting or using cuttings of corals to improve the number of outplants, and/or moving forward to other locations.

Nursery-grown corals, fragments of opportunity of multiple species, as well as micro-fragments and cuttings are directly outplanted to the reef using Portland marine cement mixed with lime to neutralize pH. Cable ties and masonry nails are also used in the case of *A. cervicornis*. An outplanting schedule with a density of one individual per square meter of reef for *A. cervicornis* and of one colony per four square meters for other species is often followed. The total spatial extent intervened through the previous stages of the project was 6 ha, but many of these corals were lost during the 2017 hurricanes. The projected spatial extent of reef rehabilitation by year 2022 in total will be 8.4 ha, with a potential to increase the area intervened to 11.7 ha depending on funding and on community-based volunteer support. The funds projected towards restoration for the period of 2019 to 2022 are $1,327,206 (2018 USD), resulting in $158,189 USD per restored ha per year or an estimated investment of $50.26 USD per coral colony. These figures are based on the direct funds spent without accounting for in-kind contributions from the community. The real total estimated budget (including community-based in-kind support) for the period of 2019 to 2022 is $2,311,280 (2018 USD) resulting in a total annual expenditure of $275,480 (2018 USD) $ha^{-1}$ or a total estimated expense of $87.53 per coral colony. The best guess of the historical (2003-2017) estimated feasibility is 0.9 (minimum of 0.5 and maximum of 1.0) based on the 6 specific project objectives. The first 14 year-long phase of the project was financed by multiple sources, including US Federal funding and minor funding from private sources. It also involved extensive volunteer work, even from SAM's staff. The conservative estimate is that approximately 80% of the work conducted between 2003 and 2017 was constituted by in-kind donations from project staff and community-based volunteers, particularly for the first 8 years of the project. There was also a large focus on community-based outreach, education, empowerment and hands on training of local community members such as students, divers and fishermen to maintain the coral nurseries and continue to outplant coral fragments to the reefs. Trained volunteer participation has exceeded 600 persons though the history of the project.

The best approximation of the current projected short-term (2019-2022) feasibility is 0.7 (minimum of 0.5 and maximum of 0.9) based on the six specific project objectives. The first 4-year sub-project of the post-hurricane long-term phase of the project will be financed through multiple sources, as described above. It will also involve extensive volunteer work, through a combination of strategies involving students, fishermen, NGOs, and an internship program. SAM also plans to involve the hospitality sector. There will also be a large focus on a combination of outreach, educational and hands on strategies to prepare the next generation of coral farmers and coral reef restoration researchers in Puerto Rico.

*Planned work*

Country: Colombia

Location: Gorgona National Natural Park, Colombian Pacific

Organization: La Fundación para la Investigación y Conservación Biológica Marina ECOMARES (Universidad del Valle, Universidad Javeriana de Cali, and Gorgona National Natural Park)

Since 2015, studies have been carried out within Gorgona National Natural Park, an island located in the Pacific coast of Colombia, at approximately 28 km from the mainland (2.9694444° N, -78.18472222° W). At present, Pacific Colombian coral reefs are in good condition, with only a few signs of degradation, therefore there is no need for coral restoration yet. Acknowledging that Gorgona's reef are going to have the same fate as all other reefs suffering from climate change, overfishing, pollution, coastal development, bleaching and diseases, in 2015 a coalition of different institutions was built to gather scientific information on coral restoration. The coalition is motivated by experimental reasons to improve restoration approaches for their use at Gorgona National Natural Park and explore sites that would be best suited for future restoration. Coral reefs in the area are built and dominated by the branching coral *Pocillopora damicornis* and 90% of the studies have focused on this species. The other species studied is the massive coral *Pavona clavus*, which is usually found in deeper areas. The patch reefs of Gorgona hold a high biodiversity and abundance of fish and benthic invertebrates, although the corals are quite dispersed and isolated from other colonies and reefs (Pizarro V., personal observation). Some of the objectives of the studies have been: 1) to determine the feasibility of coral nurseries in the area; 2) to determine the minimum coral fragment size of *P. damicornis* for successful survival and growth in a coral nursery; 3) to find the optimal fragment size for outplanting in terms of survival and coral growth; 4) to determine the effect of fish predation on *P. damicornis* during the outplanting; and 5) to evaluate the use of enriched substrates for the massive coral species *P. clavus*. So far, the duration for each study has been on average 1.5 years. However, this group expects to have projects running over the next 3 – 5 years depending on the coral species: three branching coral species and five massive coral species for future coral reef restoration. The group's expertise in outplanting has been focused towards *P. damicornis*. For this coral species, Portland cement mixed with sand and freshwater was used. So far, no information is available to determine the spatial extent (area) of restored habitat that will be obtained. The cost for running the projects have been lower than expected because they are mostly experimental and have not carried out formal coral reef restoration activities yet. In 2018 the budget was $10,000 USD. Preliminary results have shown that coral restoration at Gorgona National Natural Park are feasible (best guess of 0.7 with minimum of 0.5 and maximum of 0.9) and success can be achieved in only a few years (i.e., 3 – 5) due to the rapid growth of *P. damicornis*.

Country: Mexico

Location: Cozumel National Natural Park & Mexican Caribbean

Organization: The Iberostar Group and CINVESTAV Group: Ecology and Coral Reef Ecosystems Laboratory

Cozumel Island (20.314565 N, -87.030132 W) is part of the Mesoamerican Reef System, the second largest barrier reef in the world, at over 1,000 km from Mexico to northern Honduras. Cozumel is located 18 km from the east coast of the Yucatan Peninsula in the northwest of the Caribbean and is 46 km long and 16 km wide. Cozumel is surrounded by 11,987 hectares of coral reef which is part of the Cozumel National Natural Park. Annual monitoring is carried out within this reef with data available from 2004 until the present. Biological monitoring has identified six reef types between a depth of 10 m and 15 m. Some of those reefs are under local threat by increased tourism on the island. Both the local coral reefs as well as the species have been monitored, which is crucial for the implementation of coral reef restoration programs. Although the Healthy Reefs for Healthy People

Initiative declared the reefs of Cozumel to be in a healthy state [50], since December 2017, Scleractinian Coral Tissue Loss Disease (SCTD) has been encountered on the reef as one of the main threats to the reefs in the Cozumel and Mexican Caribbean region. The Iberostar Group and CINVESTAV Group have recently started a collaboration with the Cozumel National Natural Park and the Mexican Secretariat of Environment and Natural Resources (SEMARNAT) to implement a comprehensive coral reef restoration program for the island and other potential sites in the Mexican Caribbean, there is no estimated duration of the restoration project, as it is still being developed under Iberostar's *Wave of Change* movement. The program will engage with local communities, universities, government entities and tourism service providers to gather sustained funding into the future. Coral reef restoration envisioned by both groups is mainly motivated by experimental, biotic (i.e., enhance biodiversity, ecosystem connectivity, and ecological resilience), idealistic and pragmatic reasons (i.e., enhanced water quality and ecosystem services, shallow-water essential fish habitat, restore depleted fisheries, enhanced tourism, and coastal protection of local coral reefs. The collaborative restoration project follows four specific objectives: 1) to develop genotyped coral nurseries, which represent the coral diversity at Cozumel Island; 2) to establish sufficient material in the coral nurseries to develop activities for education, research, technological innovation, recreation and tourism; 3) to yield sufficient material for the establishment of transplant zones; and 4) to collect gametes during the spawning season for larval rearing and use the larval propagation technique to grow sexual recruits at the transplantation site. The recently established coral restoration group between Iberostar and CINVESTAV aims to start with the development of four genotyped coral nurseries, two for *Acropora palmata* (3 and 5 m water depth) and two for *A. cervicornis* (10 and 13 m water depth). Each nursery will have 5 structures with a carrying capacity of approximately 40 fragments each enabling growth of 800 corals at a time. The short-term (within one year) goal of the project is to gather enough genetic material for the development of activities related to education, research, technological innovation, recreation and tourism. In the medium term (within three years), these nurseries will provide the necessary genetic material to establish outplanting sites. The project envisions gamete collection during the spawning season to reseed the transplant site with laboratory-grown coral larvae. The project intends to create new nurseries in the future and incorporate additional reef-building coral species such as *Pseudodiploria* spp., *Siderastrea* spp., *Diploria labyrinthiformis* and *Orbicella* spp. The group is open to new partners interested in participating in the project. This program will not only be important and necessary to achieve the conservation of Critically Endangered coral species and recover the ecosystem functions and services provided by the reefs on Cozumel, but it will also become a platform for environmental research and education for the area. It is envisioned for the project to become a major tourist attraction on the island, with a scientific basis, which can serve as a buffer for adjacent reef areas.

### *Tables*

**Table S1:** Overview of coral restoration techniques employed by the projects in the Caribbean and Eastern Tropical Pacific.

| Country, Location, Organization | Direct transplantation | Coral gardening | Micro-fragmentation | Larval propagation |
|---|---|---|---|---|
| **Implemented and in progress as of 2019** | | | | |
| Colombia, Taganga, Caribbean Sea, Alianza Coralina Taganga | | | X | |
| Colombia, San Andres and Providencia Islands, Caribbean Sea, Corales de Paz | | X | X | |
| Costa Rica, Golfo Dulce, Eastern Tropical Pacific, Raising Coral Costa Rica | | X | X | |
| Dominican Republic, Bayahibe, Caribbean Sea, FUNDEMAR | | X | | X |
| Dominican Republic | | X | | |

| | | | | |
|---|---|---|---|---|
| Location, Bayahibe, Caribbean Sea, The Iberostar Group | | | | |
| Dominican Republic, Punta Cana, Caribbean Sea, Fundación Grupo Puntacana | | X | X | |
| Mexico, Chetumal Stakeholder, Caribbean Sea, Oceanus A.C. | | X | | |
| Mexico, Mexican Caribbean, Caribbean Sea, CORALIUM, Universidad Nacional Autónoma de México | | | | X |
| Puerto Rico, Culebra Island, Caribbean Sea, Sociedad Ambiente Marino | X | X | X | |
| **Planned work** | | | | |
| Colombia, Gorgona National Natural Park, Eastern Tropical Pacific, ECOMARES | | X | | |
| Mexico, Cozumel National Natural Park, Caribbean Sea, The Iberostar & CINVESTAV Group: | | X | | |

**Table S2:** List of *in situ Acropora cervicornis* propagation nurseries by FUNDEMAR. Abbreviation: *Acropora cervicornis* (Acer).

| **Location** | **Site** | **Latitude** | **Longitude** | **Year established** | **Species** |
|---|---|---|---|---|---|
| Coral reefs of the south-eastern zone (South of marine sanctuary) | FUNDEMAR | 18.3609° | -68.84515° | 2011 | Acer |
| | CATALONIA | 18.34029° | -68.82735° | 2014 | Acer |
| | DREAMS | 18.36965° | -68.85346° | 2015 | Acer |
| | SCUBA FUN | 18.34436° | -68.83389° | 2015 | Acer |
| | VIVA | 18.34590° | -68.83274° | 2015 | Acer |
| | IBEROSTAR | 18.33915° | -68.82641° | 2016 | Acer |
| | CANOA | 18.34151° | -68.82789° | 2016 | Acer |
| | CATUANO | 18.19426° | -68.78007° | 2018 | Acer |
| Coral reefs of the south-eastern zone (North of marine sanctuary) | CATALONIA - BÁVARO | 18.65728° | -68.35378° | 2016 | Acer |
| Las Terrenas | Coralas Las Terrenas Foundation | 19.33657° | -69.57264° | 2017 | Acer |

**Table S3:** List of sites where asexual colonies of *Acropora cervicornis* propagated in the nurseries have been outplanted to date by FUNDEMAR.

| Location | Site | Latitude | Longitude | Outplanting year |
|---|---|---|---|---|
| Coral reefs of the south-eastern part of the marine sanctuary | Magallán | 18.3609° | -68.84515° | 2013-2017 |
| | Coralina | 18.37083° | -68.84840° | 2013 |
| | Pepito I | 18.34533° | -68.83232° | 2014-2016 |
| | Pepito II | 18.34424° | -68.83087° | 2014-2016 |
| | Atlantic Princess | 18.3691° | -68.85225° | 2016-2019 |
| | Costa Romántica | 18.38245° | -68.85043° | 2016 |
| | Playita | 18.37308° | -68.85326° | 2017-2018 |
| | Vivero Catalonia Bávaro | 18.65728° | -68.35378° | 2018 |
| | Vivero Scuba Fun | 18.34436° | -68.83389° | 2018 |
| | Vivero Fundemar | 18.3609° | -68.84515° | 2019 |
| | Vivero Iberostar | 18.33915° | -68.82641° | 2019 |
| | Vivero Catalonia | 18.34029° | -68.82735° | 2019 |

**Table S4:** List of monitoring spawning sites of FUNDEMAR. Abbreviation: *Acropora cervicornis* (Acer), *Colpophylia natans* (Cnat), *Montastraea cavernosa* (OCav), *Orbicella annularis* (Oann), *Orbicella faveolata* (Ofav), *Dendrogyra cylindrus* (Dcyl), and *Diploria labyrinthiformis.*

| Location | Site | Latitude | Longitude | Sight year | Species |
|---|---|---|---|---|---|
| Coral reefs of the south-eastern part of the marine sanctuary | Vivero Fundemar | 18.3609° | -68.84515° | 2015-2018 | Acer<br>Cnat<br>Ocav<br>Oann<br>Ofav |
| | Playita | 18.37308° | -68.85326° | 2017-2018 | Acer<br>Dcyl<br>Dlab |

**Table S5:** Estimated feasibility of the reef restoration major objectives of the Iberostar Group.

| Objective | Estimated feasibility | Minimum feasibility | Maximum feasibility |
|---|---|---|---|
| 1. Determine current intraspecific diversity | 0.4 | 0.2 | 0.8 |
| 2. Enhance intra- and inter-specific diversity | 0.6 | 0.3 | 0.8 |
| 3. Maintain *in situ* and *ex situ* genetic bank | 0.6 | 0.2 | 1 |
| 4. Engage hotel clients and staff | 0.6 | 0.3 | 0.8 |
| 5. Characterise individual physiological traits of corals | 0.4 | 0.1 | 0.7 |
| 6. Enhance resilience in restored reefs. | 0.4 | 0.1 | 0.8 |
| **Mean feasibility** | **0.5** | **0.2** | **0.8** |

**Table S6:** List of all restoration sites of Oceanus A.C.

| Location | Site | Latitude | Longitude |
|---|---|---|---|
| Punta Allen | Punta Allen | -87.418218 | 19.739874 |
| Chacmool | Maria Elena La poza | -87.441697 | 19.457694 |
| Herrero | Transplante Faro | -87.43832 | 19.31689 |
| Veracruz | Arrecife Anegada de Adentro PNSAV | -96.061416 | 19.230431 |
| Veracruz | Arrecife Pájaros PNSAV | -96.081941 | 19.186026 |
| Xcalac | La Poza XC | -87.82558 | 18.260935 |
| Xcalac | Parche Chol | -87.831978 | 18.216738 |
| Puerto Morelos | Rodman | -86.853884 | 20.870556 |
| Puerto Morelos | Jardines | -86.880334 | 20.831163 |
| Puerto Morelos | La Pared | -86.876528 | 20.823953 |
| Playa del Carmen | Mayakobá | -87.015281 | 20.678092 |
| Mahahual | Rio Bermejo | -87.715526 | 18.685029 |
| Mahahual | Margarita del Sol | -87.71932 | 18.667874 |
| Xcalac | Acocote | -87.807633 | 18.341611 |
| Cozumel | Chankanaab | -86.995572 | 20.442498 |
| Cozumel | El Palmar | -86.986468 | 20.459119 |

**Table S7:** List of sites where coral sexual recruits produced in the laboratory have been outplanted by CORALIUM. Abbreviations: *Acropora palmata* (Apal), *Orbicella faveolata* (Ofav), *Orbicella annularis* (Oann), *Pseudodiploria strigosa* (Pstr), and *Diploria labyrinthiformis* (Dlab).

| Location | Site | Latitude | Longitude | Outplanting year | Species |
|---|---|---|---|---|---|
| Cancún | Cuevones | 21.161694° | -86.740972° | 2015-2016 | Apal |
| Puerto Morelos | Cuevones | 20.91897° | -86.830149° | 2017 | Apal |
| | Manchones | 20.97855° | -86.800631° | 2018 | Ofav<br>Oann |
| | Picudas | 20.88385° | -86.848144° | 2015-2017 | Apal |
| | Jardines | 20.830981° | -86.874981° | 2018 | Apal<br>Oann<br>Ofav<br>Pstr<br>Dlab |
| Punta Allen | Pajaritos | 19.6722861° | -87.417244° | 2018 | Apal<br>Ofav<br>Dlab |
| Puerto Herrero | Quebrado | 19.365458° | -87.439407° | 2015 | Apal |
| Xcalak | Portillas | 19.6722861° | -87.833233° | 2015 | Apal |

**Supplementary Excel file**

**Table S8:** Spatial information of all restoration projects (including sites of nurseries and outplanting sites).

File: Data_coral_reef_restoration.xlsx

Tab: 'Spatial information'

**Table S9:** Motivations or rationales for conducting the coral reef restoration projects grouped by the categories: 1) biotic; 2) experimental; 3) idealistic; 4) legislative; and 5) pragmatic.

File: Data_coral_reef_restoration.xlsx

Tab: 'Motivations'

**Table S10:** Specific objectives of all coral reef restoration projects grouped into the categories: 1) enhance ecosystem services for the future; 2) optimize/scale-up restoration approach; 3) promote coral reefs; 4) conservation stewardship; 5) provide alternative, sustainable livelihood opportunities; 6) reduce population declines and ecosystem degradation; and 7) re-establish a self-sustaining, functioning reef ecosystem.

File: Data_coral_reef_restoration.xlsx

Tab: 'Specific objectives'

**Table S11:** Cost, spatial extent of intervention, project duration and feasibility of coral reef restoration projects.

File: Data_coral_reef_restoration.xlsx

Tab: 'Cost & feasibility'
